# Submillimeter diffusion MRI using an in-plane segmented 3D multi-slab acquisition and denoiser-regularized reconstruction

**DOI:** 10.1101/2024.10.10.617536

**Authors:** Ziyu Li, Silei Zhu, Karla L. Miller, Wenchuan Wu

## Abstract

High-resolution diffusion MRI (dMRI) provides valuable insights into brain microstructure, particularly at submillimeter resolutions, where it enables more precise delineations of curved and crossing white matter pathways. However, achieving high-quality submillimeter dMRI in-vivo poses significant challenges due to the intrinsically low signal-to-noise ratio (SNR), along with the long echo spacing, readout time, and TE required for the large matrix size, leading to significant image distortion, T2* blurring, and T2 signal decay. In this study, we propose a novel acquisition and reconstruction framework to overcome these challenges. Based on numerical simulations, we introduce an in-plane segmented 3D multi-slab acquisition that leverages the optimal SNR efficiency of 3D multi-slab imaging while reducing echo spacing, readout times, and TE using in-plane segmentation. This approach minimizes distortion, improves image sharpness, and enhances SNR. Additionally, we develop a denoiser-regularized reconstruction to suppress noise while maintaining data fidelity, which reconstructs high-SNR images without introducing substantial blurring or bias. Comprehensive in-vivo experiments demonstrate that our method consistently produces high-quality dMRI data at 0.65 mm and 0.53 mm isotropic resolutions on a 3T scanner. The submillimeter dMRI datasets reveal richer microstructural details, reduce gyral bias, and improve U-fiber mapping compared to prospectively acquired 1.22 mm diffusion data. Our method demonstrates robustness at 7T and generates high-SNR 0.61 mm diffusion datasets, showing excellent agreement with previous post-mortem studies at the same scanner. Implemented using the open-source, scanner-agnostic framework Pulseq, our approach may facilitate broader adoption across different scanner platforms to benefit a wider range of applications. These results underscore the potential of our method to advance medical image analysis and neuroscientific research on human brain connectivity.

## 1. Introduction

High-resolution diffusion MRI (dMRI) offers a powerful tool for investigating detailed brain microstructure. The benefits of submillimeter dMRI have been demonstrated in post-mortem studies (Miller et al., 2011; Tendler et al., 2022), where it provides more precise delineations of curved and crossing white matter pathways (e.g., transverse pontine fibers) compared to conventional resolutions (e.g., 2 mm). Additionally, submillimeter dMRI may help address a known challenge in dMRI fiber tracking (tractography) called “gyral bias”, where tracked fibers tend to terminate at gyral crowns instead of accurately following the gyral walls (Cottaar et al., 2021; Zhu et al., 2024). Higher spatial resolutions are also advantageous for the identification of short cortical association fibers, commonly known as U-fibers, which connect cortical regions between adjacent gyri (Song et al., 2014). This is particularly important for studying the brain’s structural connectivity (Ouyang et al., 2017). Furthermore, high-resolution dMRI holds great promise for precisely delineating small but crucial subcortical structures, which can serve as targets for deep brain stimulation in treating conditions such as Parkinson’s disease, essential tremor, and dystonia (Maffei et al., 2022; Saleem et al., 2021).

Despite these appealing advantages, achieving high-resolution in-vivo dMRI, especially at submillimeter levels, presents considerable challenges. For the conventional 2D single-shot echo-planar imaging (EPI), a major hurdle is the intrinsically limited SNR due to the small voxel size and the long TE associated with the large number of phase encoding lines. The long TR necessary to cover a large number of slices reduces SNR efficiency, which is optimized with TR=1-2s for spin echo-based dMRI (Frost et al., 2014). Additionally, the large imaging matrix size also results in extended echo spacing and readout times, leading to significant image distortion that compromises image anatomical fidelity, and T2* blurring that affects effective resolution (Feizollah & Tardif, 2023). Moreover, the challenges in delivering sharp slice-selective RF pulses to resolve thin slices limit the achievable through-plane resolution.

To overcome these challenges, previous studies have proposed super-resolution-based approaches that have shown potential in enabling submillimeter dMRI. One notable technique is gSlider (Liao et al., 2021; Liao et al., 2023; Setsompop et al., 2018), which uses 2D acquisitions to excite a thin slab multiple times with distinct, orthogonal RF pulses, allowing thin slices to be resolved from the slab (typically 5 slices per slab). Coupled with the simultaneous multi-slice (SMS) technique, gSlider-SMS can effectively shorten the TR and improve SNR efficiency. More recently, super-resolution-based submillimeter dMRI by acquiring multiple thick-slice volumes with rotated field-of-view (FOV) to resolve a high-resolution volume has also been proposed (Dong et al., 2024). However, a potential drawback of these super-resolution methods is the blurring effect that can result from the use of conditioning techniques (e.g., Tikhonov regularization) when solving the inverse problem. Furthermore, despite achieving shorter TRs than conventional 2D EPI, their TRs (typically longer than 3 seconds) still do not fall within the optimal range for SNR efficiency.

An alternative and promising approach for achieving high-resolution, SNR-efficient dMRI is 3D multi-slab imaging, which is compatible with shorter TRs of 1-2 seconds (Engström & Skare, 2013; Frost et al., 2014; Li et al., 2023; Wu, Poser, et al., 2016). This technique divides the whole brain into several slabs, each with a thickness of less than 2 cm to ensure that motion-induced phase variations can be effectively captured by a 2D navigator (Engström & Skare, 2013). Within each slab, 3D EPI is typically employed for encoding, offering superior SNR and image sharpness due to the use of orthogonal Fourier bases. However, conventional 3D multi-slab imaging often uses a single shot to cover one kz plane, resulting in extended TE and readout times, which can lead to significant SNR penalties and T2* blurring, particularly at submillimeter resolutions.

In this study, we present a novel acquisition and reconstruction framework designed to achieve submillimeter in-vivo dMRI. We leverage 3D multi-slab imaging for superior SNR efficiency, with in-plane segmented EPI used to shorten echo spacing, readout times, and TE, thereby reducing distortion, T2* blurring, and improving SNR. A denoiser-regularized reconstruction approach is proposed to effectively suppress noise during reconstruction while maintaining consistency with the acquired data to avoid biases. In-vivo experiments at 3T demonstrate that our method produces high-quality submillimeter dMRI at 0.65 mm and 0.53 mm isotropic resolutions. The efficacy of our approach is further validated at a higher field strength of 7T, where it consistently delivers high-quality data. Comprehensive diffusion analyses of the acquired submillimeter datasets reveal significantly richer microstructural details, underscoring our method’s potential to address the “gyral bias” and improve U-fiber tracking compared to conventional 1.22 mm resolution data. These findings hold considerable promise for advancing neuroscientific research into the human brain.

## 2. Methods

### 2.1. In-plane Segmented 3D Multi-slab Acquisition

In conventional 3D multi-slab diffusion imaging, each shot typically covers a kz plane using a single EPI readout for efficient data acquisition (Dai et al., 2021; Engström & Skare, 2013; Wu, Poser, et al., 2016). However, for high-resolution EPI with a large matrix size, this single-shot method can result in extended echo spacing, readout durations, and TE, leading to significant image distortion, T2* blurring, and reduced SNR.

In-plane segmented multi-shot acquisitions hold promise in mitigating these issues. Previous studies have proposed readout-segmented 3D multi-slab EPI (Frost et al., 2014; Wu, Koopmans, et al., 2016), but these methods have not demonstrated submillimeter-level resolution, likely due to limitations in maximum slew rate. An alternative approach is phase-encoding segmented EPI, where each shot undergoes a high under-sampling factor along the phase encoding direction (ky). This effectively shortens the effective echo spacing, readout duration, and TE. The effective echo spacing and associated image distortion decreases inversely proportional to the number of in-plane segments (*N*_*seg*_). For the effective resolution and SNR, we quantitatively assess how they are affected by *N*_*seg*_ at 0.6 mm and 1 mm resolutions, assuming the data are fully sampled with effective acceleration factor *R*_*eff*_ = *N*_*seg*_/*N*_*acq*_ = 1. We extend the single-shot simulation framework by Feizollah and Tardif (Feizollah & Tardif, 2023) for multi-segment acquisitions, modifying the parameters to reflect realistic 3D multi-slab dMRI acquisitions: TR=2.5s, maximum gradient amplitude *G*_*max*_=80 mT/m, RF excitation duration=6 ms, RF refocusing duration=10 ms, b-value=1000 s/mm^2^, and bandwidths of 992 Hz and 1384 Hz for 0.6 mm and 1 mm acquisitions, respectively. The T1, T2, and T2* values are set to 800/79.6/53.2ms at 3T and 1200/47/26.8ms at 7T for white matter (Feizollah & Tardif, 2023).

The effective resolution is quantified by the full-width-half-maximum (FWHM) of the point spread functions simulated under various acquisition parameters. For the partial Fourier (PF) sampling, conjugate symmetric filling and zero-padding are investigated. The SNR is quantified based on:

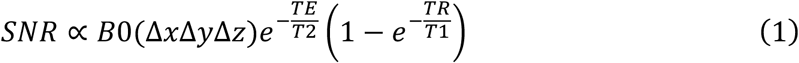

where B0 is the main magnetic field strength, Δ*x*, Δ*y*, and Δ*z* are the effective resolution along readout, phase-encoding, and slice-selection directions, respectively, and (Δ*x*Δ*y*Δ*z*) is the effective voxel size. While Δ*y* is determined by FWHM, Δ*x* and Δ*z* are assumed to match the nominal resolution. The TE is simulated under different acquisition conditions.

As detailed in the Results section, the simulations indicate that a large *N*_*seg*_ is necessary for improving effective resolution and SNR. Based on our simulation results, we select *N*_*seg*_ of 6 for 3T and 8 for 7T acquisitions of submillimeter diffusion data, by considering the balance between scan time, effective resolution, and SNR.

For the sampling order of in-planed segmented 3D EPI, we choose to acquire all in-plane ky segments of a given kz in immediate succession before moving to the next kz (“ky-kz” in Supplementary Fig. 1a). Compared to the alternative order where all kz planes for one ky segment are acquired before proceeding to the next segment (“kz-ky” in Supplementary Fig. 1b), our chosen sampling order reduces the inconsistencies between in-plane segments, improving motion robustness and reducing image artifacts (Ivanov et al., 2015; Polimeni et al., 2016), especially for the b=0 image acquisition due to the pronounced cerebrospinal fluid (CSF) signal aliasing introduced by inter-segment inconsistency (Supplementary Fig. 1).

### 2.2. Denoiser-regularized Reconstruction

The accuracy of high-resolution dMRI reconstruction is often limited by the inherently low SNR. Image denoising is a common strategy to enhance SNR, with various methods available both for general natural images (e.g., Non-Local Means (Buades et al., 2011), BM3D (Dabov et al., 2007), BM4D (Maggioni et al., 2012) and specifically for dMRI (e.g., MPPCA (Veraart et al., 2016), NORDIC (Moeller et al., 2021)). However, these methods typically function as standalone post-processing steps, which can introduce biases and blurring during dMRI denoising (Manzano Patron et al., 2024).

Our approach integrates denoising within a SPIRiT-based (Lustig & Pauly, 2010) multi-shot reconstruction framework, aiming to suppress noise while minimizing biases and blurring by enforcing data consistency:

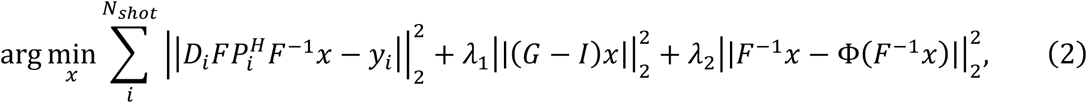

where *x* is the desired fully-sample, high-SNR, multi-coil k-space data to be reconstructed, *N*_*shot*_ is the total number of shots, *D*_*i*_ is the shot-sampling mask for the *i*^*th*^ shot, *F* is the Fourier Transform, *P*_*i*_^*H*^is the conjugate of 2D navigator phase for addressing motion-induced phase variance (Engström & Skare, 2013; Miller & Pauly, 2003) for the *i*^*th*^ shot, *y*_*i*_ is the acquired data for the *i*^*th*^ shot, *G* is the SPIRiT kernel trained on coil calibration data, *I* is the identity matrix, Φ is the denoiser, and *λ*_1_, *λ*_2_ are the weights for SPIRiT and denoiser regularizations, respectively.

Directly solving Eq. 2 is challenging due to the computational complexity of differentiating Φ, especially for modern sophisticated denoisers. We leverage the “plug-and-play” approach (Ahmad et al., 2020), iteratively alternating image denoising with forward-model-based reconstruction. At the *k*^*th*^ iteration:

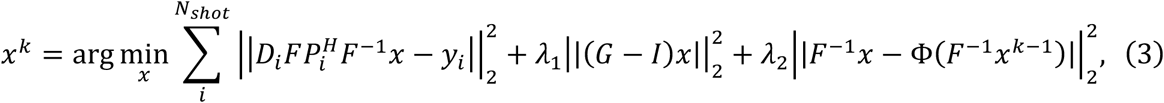

which enables the incorporation of advanced denoisers while enforcing the data consistency by minimizing 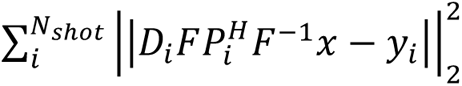 and the consistency with the calibration data by minimizing the SPIRiT constraint 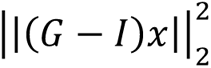. The proposed denoiser-regularized SPIRiT reconstruction is referred to as “DnSPIRiT” hereafter.

### 2.3. In-vivo Experiments

We adapted a 3D multi-slab spin-echo diffusion MRI sequence (Wu, Poser, et al., 2016) to incorporate in-plane segmented acquisitions following the sampling order illustrated in Supplementary Fig. 1a. The sequence was implemented on Siemens platform as well as an open-source, scanner agnostic framework “Pulseq” (Layton et al., 2017) to promote broader accessibility and applications. In-vivo experiments at 3T (Siemens Prisma, Erlangen, Germany) and 7T (Siemens Magnetom, Erlangen, Germany) were conducted, both with 32-channel receive coils. Written informed consents in accordance with local ethics were obtained from subjects before scanning. Submillimeter dMRI data at 0.65mm, 0.61mm, and 0.53mm isotropic resolutions were acquired using the following protocols, with key parameters listed in Table 1:

**Table 1.** Submillimeter dMRI acquisition parameters. a. TE1 is the imaging echo time and TE2 is the navigator echo time. b. For the 3T 0.53mm Protocol, the 3 acquired ky segments are evenly spaced. c. ESeff refers to the effective echo spacing (i.e., echo spacing/R).

| Res. (mm<sup>3</sup>) | B0 | Matrix size | PF | TE1/TE2/TR (ms)<sup>a</sup> | $N_{seg}/N_{acq}$<sup>b</sup> | ES<sub>eff</sub> (ms)<sup>c</sup> | $T_{acq}$ per dir. |
| --- | --- | --- | --- | --- | --- | --- | --- |
| 0.65 | 3T | 336×336×164 | 1 | 102/195/2500 | 6/6 | 0.21 | 6min |
| 0.53 | 3T | 414×414×186 | 3/4 | 80/201/2000 | 6/3 | 0.22 | 2.7min |
| 0.61 | 7T | 360×360×174 | 1 | 87/153/2600 | 8/8 | 0.15 | 7.6min |

#### 3T 0.65 mm Protocol

A 6-direction, fully-sampled dataset at 0.65mm isotropic resolution was acquired at 3T to validate the efficacy of the proposed acquisition and reconstruction framework. Nine slabs were acquired, each containing 20 slices, with a 20% oversampling along the kz direction to mitigate slab boundary aliasing, resulting in 24 slices per slab. Adjacent slabs were overlapped by 2 slices, leading to a total of 164 slices covering 106.6 mm along the superior-inferior direction. Slab excitation was achieved using a Shinnar-Le-Roux (SLR) pulse (Pauly et al., 1991) with a time-bandwidth product of 8 for excitation and 12 for refocusing. The pulse durations were 6 ms for excitation and 9 ms for refocusing, with flip angles set at 90° for excitation and 160° for refocusing to reduce slab boundary saturation artifacts. The navigator was acquired for each segment with a lower parallel imaging acceleration factor of *R*_*nav*_=4 for more robust reconstruction. Since the motion-induced phase is typically smooth (Chen et al., 2013), the distortion difference between the image and the navigator is not expected to significantly affect the reconstruction quality. Diffusion-weighted images were acquired along anterior-posterior phase-encoding direction with b=1000 s/mm^2^ and 6 diffusion directions with 2 b=0 image volumes along opposite phase-encoding directions for distortion correction. The total scan time was 48 min.

#### 3T 0.53 mm Protocol

To enable more advanced diffusion analyses at an ultrahigh resolution, a 20-direction dataset at 0.53mm isotropic resolution with *R*_*eff*_=2 was acquired at 3T. For this protocol, the number of slabs was 8 with 25 slices per slab, with 8% oversampling along kz applied (27 acquired slices per slab). Neighboring slabs were overlapped by 2 slices, resulting in 186 slices, providing a ∼100 mm coverage. An SLR pulse with matched parameters with the 3T 0.65mm Protocol was applied for slab excitation. The navigator was acquired with *R*_*nav*_=4. Diffusion-weighted images were acquired along anterior-posterior phase-encoding direction with b=1000 s/mm^2^ and 20 diffusion directions. Two fully-sampled b=0 image volumes along opposite phase-encoding directions were acquired. The total scan time was 65 min.

#### 7T 0.61 mm Protocol

To demonstrate the robustness of our method at higher field strength, a 6-direction dataset at 0.61mm isotropic resolution was acquired at 7T. Nine slabs, each containing 20 slices, were used, with 10% oversampling along kz (22 slices per slab). Adjacent slabs were overlapped by 2 slices, resulting in 174 slices in total. A sinc pulse with excitation and refocusing duration=6 and 8ms was used for slab excitation. The navigator was acquired with *R*_*nav*_ =6. Diffusion-weighted images were acquired along anterior-posterior phase-encoding direction with b=1000 s/mm^2^ and 6 diffusion directions and 2 b=0 images. The total scan time was 61 min.

For comparison, dMRI data at conventional resolutions of the same subjects were acquired. At 3T, a dMRI dataset at 1.22 mm isotropic resolution was obtained using conventional 3D multi-slab EPI with the following parameters: matrix size=180×180×91, R=3, PF=3/4, TE1/TE2/TR=70/150/2700 ms, number of slabs=10, slices per slab=10, kz oversampling=20%, number of overlapped slices between slabs=1, *T*_*acq*_ per volume=32.4s, 24 diffusion encoding directions (b=1000 s/mm^2^) and 4 b=0 images (with 2 images acquired along opposite phase encoding directions), total scan time=15.1 min. At 7T, another dMRI dataset at 1.05 mm isotropic resolution was acquired using the 3D multi-slab acquisition described in (Li et al., 2023) with matrix size=210×210×115, 48 diffusion encoding directions (b=1000 s/mm^2^) and 6 b=0 images, R=3, TE1/TE2/TR=82/150/1800 ms, *T*_*acq*_=36s per volume and ∼33 min in total. Additionally, a T1-weighted (T1w) anatomical image at 0.7 mm isotropic resolution was acquired at 3T using magnetization-prepared rapid gradient-echo imaging (MPRAGE) (Mugler III & Brookeman, 1990) following the protocol from the Human Connectome Project (HCP) (Glasser et al., 2016).

### 2.4. Reconstruction Details

The image reconstruction was conducted in MATLAB 2021a (Mathworks, Natick, MA, USA). The SPIRiT kernels *G* in Eq. 3 was trained using a whole-brain gradient echo coil calibration scan (∼2 min acquisition time). All 32-coil k-space data were compressed to 8 coils (Zhang et al., 2013).

The 2D navigator images were reconstructed with 2D GRAPPA (Griswold et al., 2002) and subsequently filtered using a k-space Hamming window of size 32×32 to reduce noise. Phase images were then extracted as estimates of motion-induced phase errors (i.e., *P*^*H*^ in Eq. 3). The reconstruction in Eq. 3 was performed for each ky-kz plane after the k-space data being first Fourier transformed along kx. The reconstructed 2D images were concatenated along the readout direction to form the whole image volume. The SPIRiT kernel size *G* was set to 5×5. Equation 3 was solved using a conjugate gradient (CG) method. The first iteration of Eq. 3 performed a standard SPIRiT reconstruction with motion-induced phase error correction.

For the 6-direction 3T 0.65 mm and 7T 0.61 mm protocols, BM4D (Maggioni et al., 2012) was employed as the denoiser, with the number of iterations (*N*_*iter*_) set to 5 (i.e., *k* ∈ [1,5]) and hyperparameters empirically selected as *λ*_1_ = 10 and *λ*_2_ = 2. During each iteration, the sum-of-squares of the reconstructed image (*F*^−1^*x*) was denoised by BM4D using adaptive noise level estimation with Rician noise distribution, while other parameters were kept at their default values. The denoised magnitude image was then multiplied by coil sensitivities, estimated from the coil calibration data using ESPIRiT (Uecker et al., 2014), to generate multi-coil complex data for the next iteration.

For the 20-direction 3T 0.53 mm protocol, NORDIC (Moeller et al., 2021) was used as the denoiser, leveraging shared information across diffusion directions. In this case, *N*_*iter*_ was set to 2, with *λ*_1_ = 10 and *λ*_2_ = 3. In each iteration, the multi-coil *F*^−1^*x* was coil-combined using the sensitivity map and then denoised by NORDIC (complex denoising), with the kernel size for g-factor map estimation set to 20×20×5, while other parameters remained at their defaults. The denoised complex image was then multiplied by the sensitivity map to generate multi-coil data for the subsequent iteration. The PF data were reconstructed using P-LORAKS (Haldar & Zhuo, 2016).

To evaluate the performance of our reconstruction method on under-sampled data, fully sampled data from the 3T 0.65 mm protocol were retrospectively under-sampled by selecting 2 and 3 evenly spaced segments from the original 6 segments, corresponding to acceleration factors of *R*_*eff*_ =3 and 2, respectively. The normalized root mean squared error (NRMSE) between the under-sampled and fully-sampled data reconstructed using SPIRiT and DnSPIRiT was calculated within a brain mask to quantify their similarities.

The 1.22 mm data acquired at 3T were reconstructed using a standard SPIRiT with motion-induced phase error correction (Li et al., 2024). A joint SPIRiT reconstruction with distortion and phase error correction detailed in (Li et al., 2023) was used for reconstructing the 1.05 mm data acquired at 7T.

### 2.5. Diffusion Analyses

Image post-processing was conducted using the FMRIB Software Library (FSL) (Jenkinson et al., 2012) unless indicated otherwise. Slab combination and correction for slab saturation artifacts were performed for the diffusion data using nonlinear inversion of slab profile encoding (NPEN) (Wu, Koopmans, et al., 2016). The images were corrected for Gibbs ringing (Bautista et al., 2021). A whole-brain field map was estimated using blip-reversed b=0 image volumes using “topup” (Andersson et al., 2003). For the 6-direction diffusion data from 3T 0.65 mm Protocol and 7T 0.61 mm Protocol, data from different diffusion directions were aligned using “eddy_correct”, and “applytopup” were used to address off-resonance distortions. For the 20-direction data from 3T 0.53 mm Protocol, the diffusion data and the field map estimated by “topup” were input to “eddy” (Andersson & Sotiropoulos, 2016) to correct for off-resonance distortions, eddy current effects, and subject motion. The 1.22 mm and 1.05 mm data were also processed by NPEN, Gibbs ringing correction, “topup”, and “eddy”, followed by “flirt” (Jenkinson & Smith, 2001) for head position alignment with high-resolution data but without upsampling. The T1w image was corrected for bias field using “fast” (Zhang et al., 2001) and co-registered to the diffusion space by applying the inverse transform obtained from “epi_reg” (Jenkinson et al., 2002; Jenkinson & Smith, 2001). Co-registered T1w image was then processed by FreeSurfer’s “recon-all” (Fischl, 2012) to produce whole-brain segmentations. All diffusion analyses were performed in the native diffusion space.

Diffusion tensor model fitting was performed on all diffusion data using “dtifit” (Smith et al., 2004). For the 20-direction 0.53 mm data and the 24-direction 1.22 mm data, fibre orientation distributions (FOD) were estimated voxel-wise using spherical deconvolution (Tournier et al., 2007; Tournier et al., 2004). A response function was first estimated with maximum harmonic degrees of 4 using MRtrix3’s “dwi2response” function and the “fa” algorithm (Tournier et al., 2019), followed by FOD estimation via MRtrix3’s “dwi2fod” function with the “csd” algorithm.

Tractography was performed to reconstruct three representative white matter bundles from the 0.53 mm and 1.22 mm datasets to evaluate the data quality. Specifically, the acoustic radiation, cingulate gyrus part of cingulum, and corticospinal tract were reconstructed with the seed, waypoint, target, and exclusion masks from “autoPtx” (Behrens et al., 2007; Behrens et al., 2003). Tractography was executed with MRtrix3’s “tckgen” function using the “iFOD2” algorithm (Tournier et al., 2010), with the cutoff value set to 0.3. For each tract, a matched number of streamlines were generated from the 0.53 mm and 1.22 mm resolution data.

To assess the “gyral bias” problem, masks were manually drawn to delineate several representative gyri on 0.53 mm and 1.22 mm data, and tractography was performed using these masks as seeds and region-of-interests (ROIs) using “tckgen” with “iFOD2”.

For U-fiber comparison between the 0.53 mm and 1.22 mm data, tractography was performed to reconstruct the short association fibers on both datasets. Supplementary Figure 2 shows the seed, waypoint, and ROI masks for the tractography. Specifically, the seed mask was set to cortical gray matter, obtained by subtracting the white matter from the cortical ribbon mask segmented from the T1w image by FreeSurfer’s “recon-all” (Supplementary Fig. 2a). The waypoint mask was the white matter mask (Supplementary Fig. 2b). The ROI mask was the cortical ribbon mask dilated using MRtrix3’s “maskfilter” with “-npass” set to four (Supplementary Fig. 2c), excluding non-relevant deep white matter tracts. The tractography was performed using “tckgen” with “iFOD2” and a minimum track length of 10 mm. On the 0.53 mm data, one seed was generated from each voxel and the cutoff value was set to 0.4. On the 1.22 mm data, two seeding mechanisms were evaluated – one seed per voxel and 12 seeds per voxel, with the latter compensating the voxel number difference due to resolution and demonstrating the impact of an increased seed number. A higher cutoff value of 0.7 was applied to reduce false positive tracts that primarily traversed in gray matter. The 0.53 mm data (one seed per voxel) and 1.22 mm data (12 seeds per voxel) produced comparable streamline numbers (1.75M and 1.84M, respectively), while the 1.22 mm data with one seed per voxel yielded 0.15M tracts.

## 3. Results

Simulation results for effective resolution and SNR across different in-plane segmentation numbers (*N*_*seg*_) for 0.6 mm and 1 mm EPI-based dMRI acquisitions at 3T are shown in Fig. 1. In all cases, T2* blurring reduces the effective resolution, but increasing *N*_*seg*_ helps mitigate this effect (Fig. 1a, c). Partial Fourier (PF) acquisitions can improve SNR by shortening the echo time (TE), though they introduce additional blurring, particularly when zero-padding (ZP) is used. In practice, phase errors lead to imperfections in conjugate symmetry (CS) in PF acquisitions, making the simple PF-CS simulation model less feasible and further degrading effective resolution. These findings emphasize the importance of using large *N*_*seg*_ to achieve higher effective resolution, especially for submillimeter imaging. Additionally, SNR improvements are also observed with increasing *N*_*seg*_due to the shorter TE (Fig. 1b, d). Notably, the SNR for 0.6 mm acquisitions is considerably lower than for 1 mm acquisitions, with over a fivefold difference (Fig. 1b, d), highlighting the substantial SNR challenges in high-resolution dMRI and the critical need for higher *N*_*seg*_ to compensate for SNR loss.

**Figure 1.**
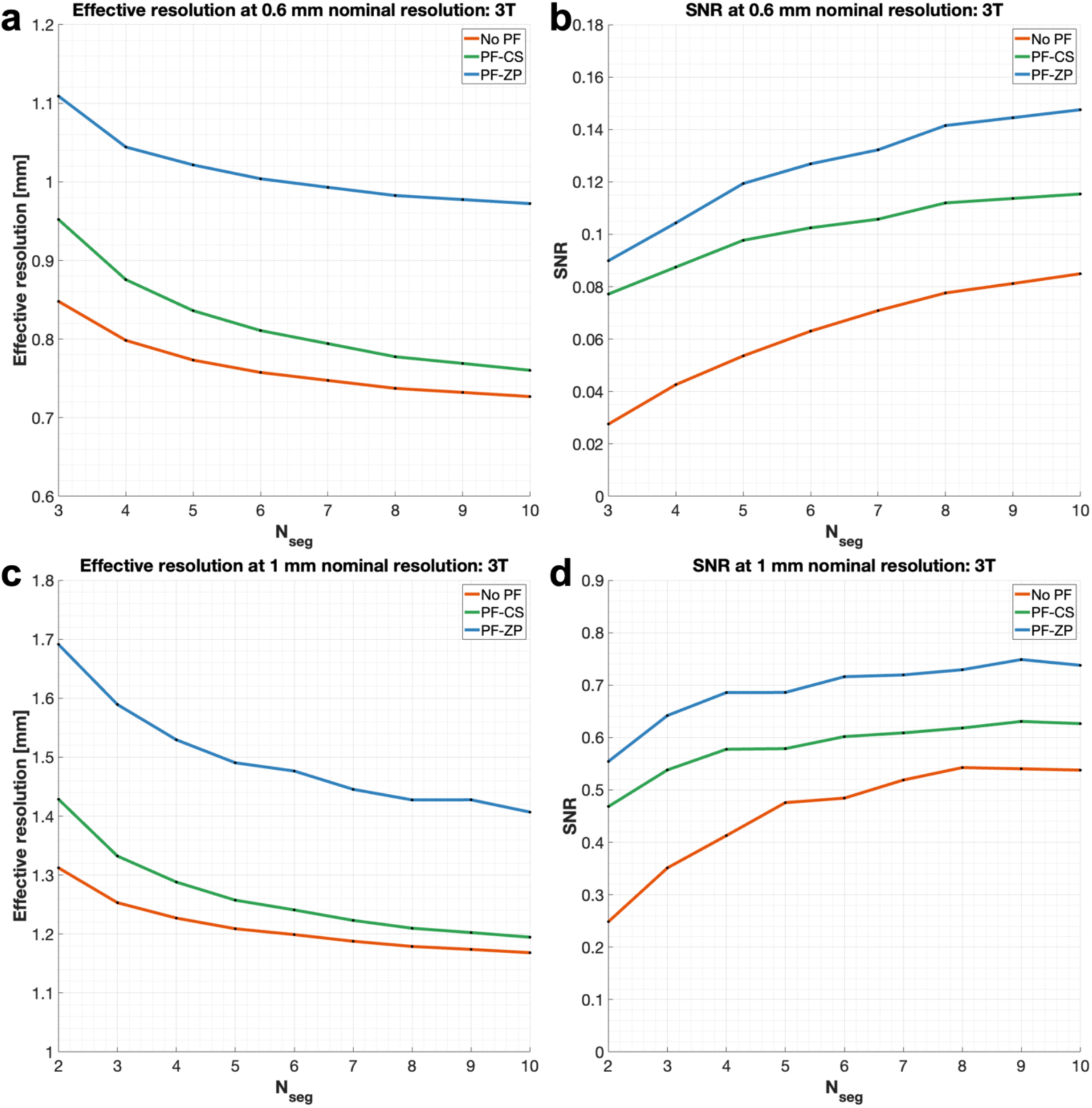
Simulation of effective resolution and SNR for high-resolution diffusion-weighted EPI at 3T. No partial Fourier (No PF, red), 6/8 PF with conjugate symmetric filling (PF-CS, green), and 6/8 PF with zero padding (PF-ZP, blue) samplings are investigated at 1mm (a, b) and 0.6mm (c, d) for white matter at 3T (TR = 2.5s, b-value = 1000 s/mm^2^, bandwidths = 992/1384 Hz/pixel for 0.6/1 mm acquisitions, T1/T2/T2* = 800/79.6/53.2ms) for different in-plane segmentation numbers (*N*_*seg*_). The SNR is calculated based on the simulated effective voxel sizes, TE, and TR.

Simulations at 7T follow similar trends (Supplementary Fig. 3). The shorter T2* at 7T intensifies T2* blurring, requiring a higher *N*_*seg*_ to reduce the readout time and achieve a resolution comparable to 3T. Moreover, the shorter T2 at 7T demands a larger *N*_*seg*_to shorten the TE and take full advantage of the SNR benefit from higher field strength. Based on these simulations, we selected *N*_*seg*_ = 6 for the 3T protocols and *N*_*seg*_ = 8 for the 7T protocol to balance scan time, effective resolution, and SNR. The simulations suggest that our three high-resolution protocols – 3T 0.65 mm (*N*_*seg*_= 6, no PF), 3T 0.53 mm (*N*_*seg*_= 6, with PF), and 7T 0.61 mm (*N*_*seg*_= 8, no PF) – achieve effective resolutions of approximately 0.8 mm.

Figure 2 demonstrates the image reconstruction results from 3T 0.65 mm Protocol. Our carefully designed acquisition scheme produces ultrahigh-resolution diffusion images with superior sharpness and adequate SNR even when using the standard SPIRiT reconstruction (Fig. 2a, b). The proposed DnSPIRiT further enhances SNR with negligible biases or blurring (Fig. 2c, d). The difference maps between SPIRiT and DnSPIRiT primarily display noise without noticeable anatomical structures or biases (Fig. 2e).

**Figure 2.**
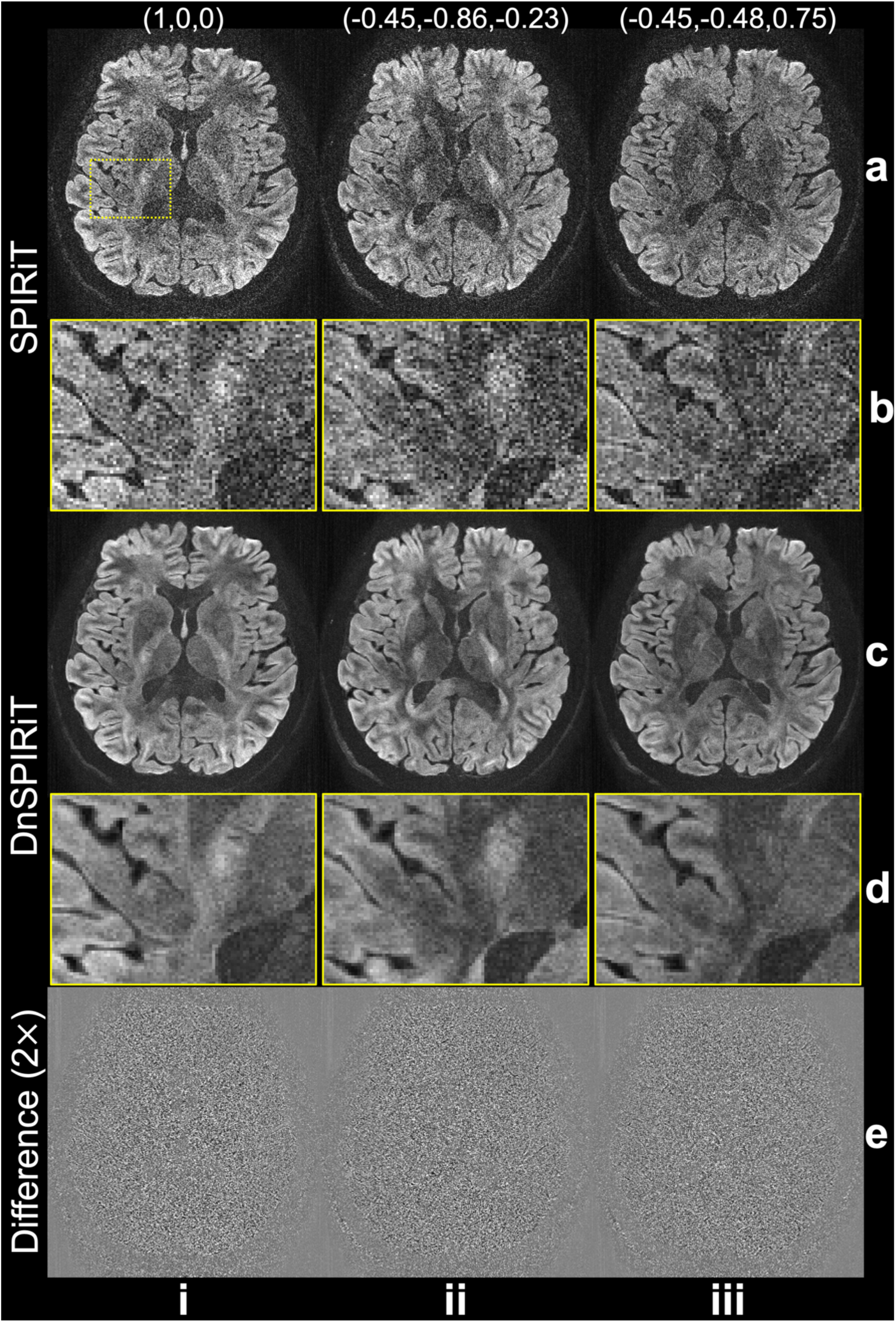
Denoiser-regularized reconstruction results. In-vivo diffusion-weighted data (b=1000 s/mm^2^) from 3T 0.65 mm Protocol along 3 representative diffusion encoding directions (i-iii, the direction is displayed above each image) are reconstructed using SPIRiT (a) and Denoiser-regularized SPIRiT (DnSPIRiT, c), with an enlarged region showing the image detail (b, d). Their difference is also shown to demonstrate the removed noise (e).

The benefit of DnSPIRiT becomes more pronounced in under-sampled reconstructions (Fig. 3). While the standard SPIRiT reconstruction exhibits significant noise for under-sampled data (Fig. 3a, b), DnSPIRiT effectively suppresses the noise (Fig. 3c, d), resulting in substantially lower NRMSE with the fully sampled reference. However, when only 2 segments are used (*R*_*eff*_=3, Fig. 3, iii), anatomical errors begin to emerge even with DnSPIRiT, presumably due to the extremely high noise level introduced by 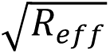 and g-factor penalty. We therefore used *R*_*eff*_=2 in 3T 0.53 mm Protocol to balance the acquisition time and reconstruction accuracy. It is worth noting that the prospective under-sampled acquisition is expected to be more robust against motion artifacts compared to retrospective under-sampling due to shortened scan time.

**Figure 3.**
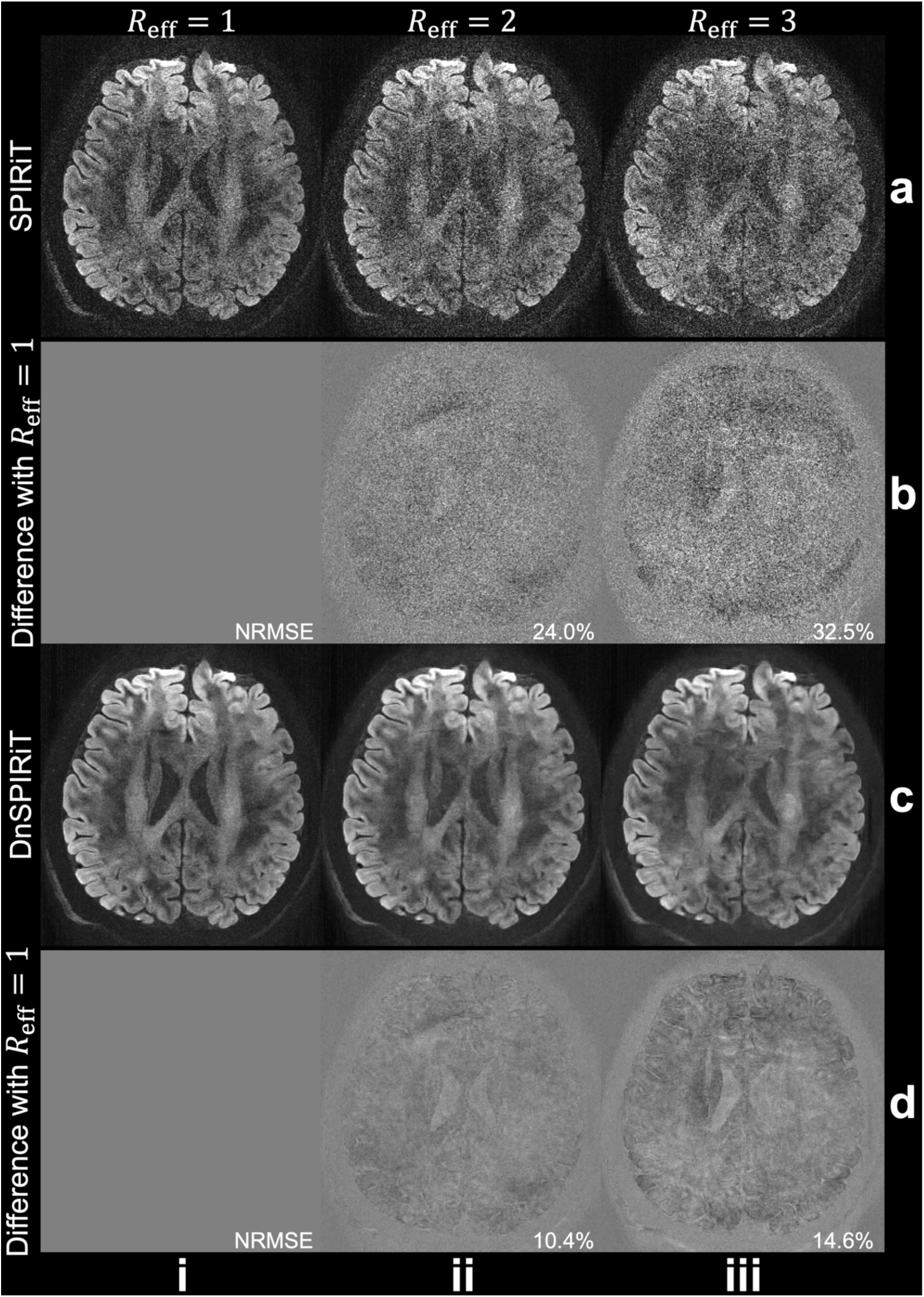
Retrospective under-sampled reconstruction. Retrospective under-sampling is applied to the fully sampled data (i) from 3T 0.65 mm Protocol by selecting 3 segments (*R*_*eff*_=2, ii) and 2 segments (*R*_*eff*_=3, iii) from the total of 6 segments. The diffusion-weighted images along (-0.45, -0.86, -0.23) reconstructed using SPIRiT (a) and DnSPIRiT (c) and their difference with fully sampled reference (b, d) are shown to demonstrate the under-sampled reconstruction fidelity. The normalized root mean squared errors (NRMSE) between the under-sampled reconstructed image and the fully sampled reference of the whole slab are calculated within a brain mask to quantify their similarity.

The whole-brain diffusion data from 3T 0.65 mm Protocol resolve rich information in brain microstructure (Fig. 4). The diffusion data exhibit sharp details and high-SNR even with only 6 diffusion encoding directions, thanks to our carefully optimized sampling and reconstruction strategies. The short effective echo spacing (0.21 ms), achieved through in-plane segmentation, significantly enhances the anatomical fidelity (Fig. 4b). For comparison, the effective echo spacing for HCP 3T diffusion imaging at 1.25 mm isotropic resolution was 0.78 ms (Uğurbil et al., 2013). Small structures such as fornix, external capsule, and anterior commissure are clearly visible on the 0.65 mm data with excellent anatomical fidelity (Fig. 4d). Moreover, the results align well with prior post-mortem data at 0.5 mm resolution (Tendler et al., 2022) (Fig. 4d).

**Figure 4.**
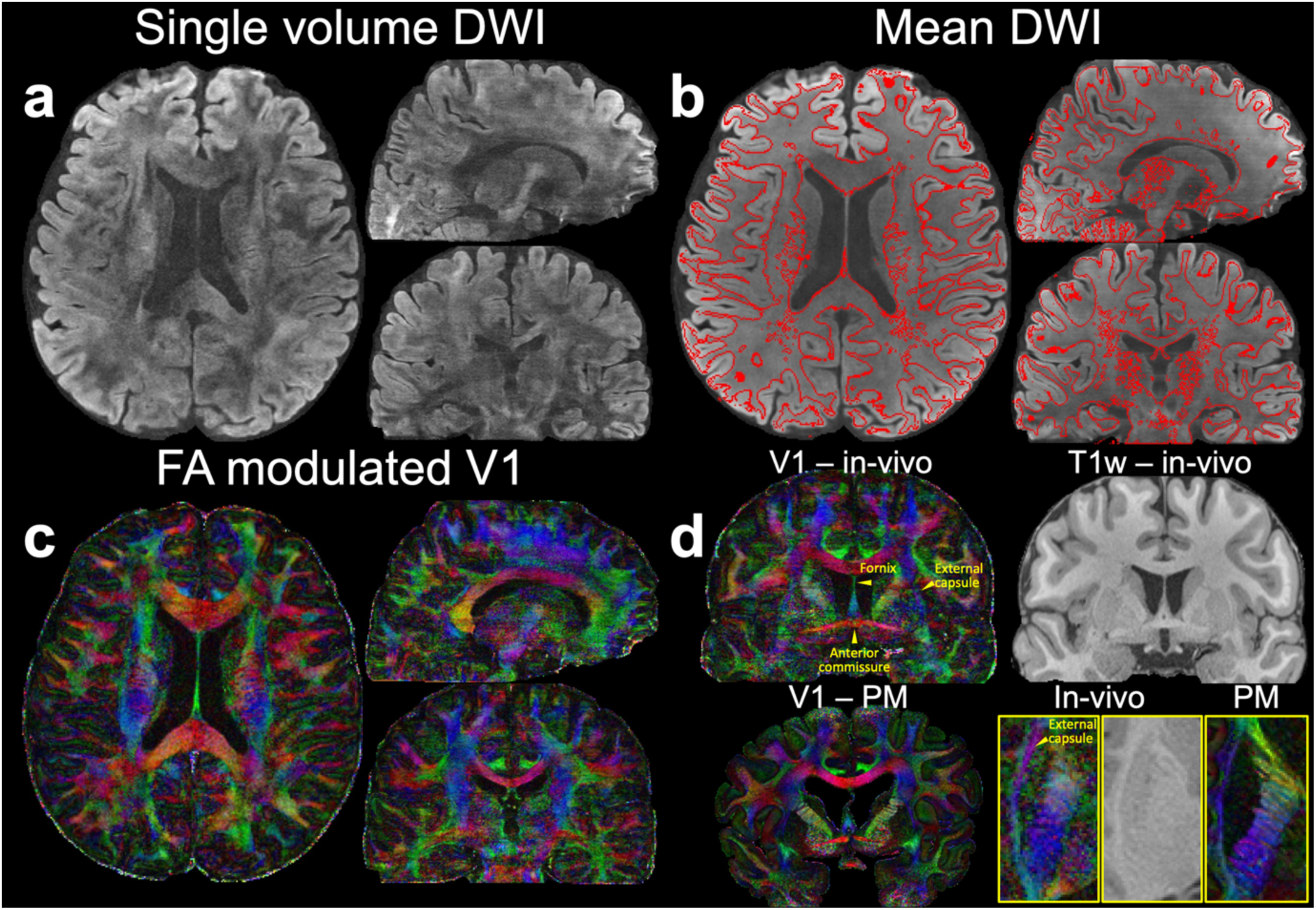
Whole-brain data from 3T 0.65 mm Protocol. Whole-brain in-vivo diffusion data (b=1000 s/mm^2^) including diffusion-weighted image (DWI) along (-0.45, 0.83, -0.32) (a), 6-direction mean DWI with gray-white matter boundary derived from the T1w image using FSL’s “fast” overlayed (b), fractional anisotropy (FA) modulated V1 (c) at 0.65 mm isotropic resolution are demonstrated. A representative sagittal view of the FA modulated V1 (V1 – in-vivo) alongside the T1-weighted image (T1w – in-vivo), highlighting key structures including the fornix, external capsule, and anterior commissure are also presented. For comparison, post-mortem data (V1 – PM) from previous studies (Tendler et al., 2022) are shown. Enlarged views of the external capsule from an axial plane are provided for both in-vivo and post-mortem data (d).

Compared to the conventional 1.22 mm resolution diffusion data, data from 3T 0.65 mm Protocol reveals substantially finer microstructure details (Fig. 5). This includes the improved visualization of the striations through the internal capsule (Fig. 5, i), the interdigitating transverse pontine fibers (Fig. 5, ii), and the clearer delineation of the cingulum bundle (Fig. 5, iii).

**Figure 5.**
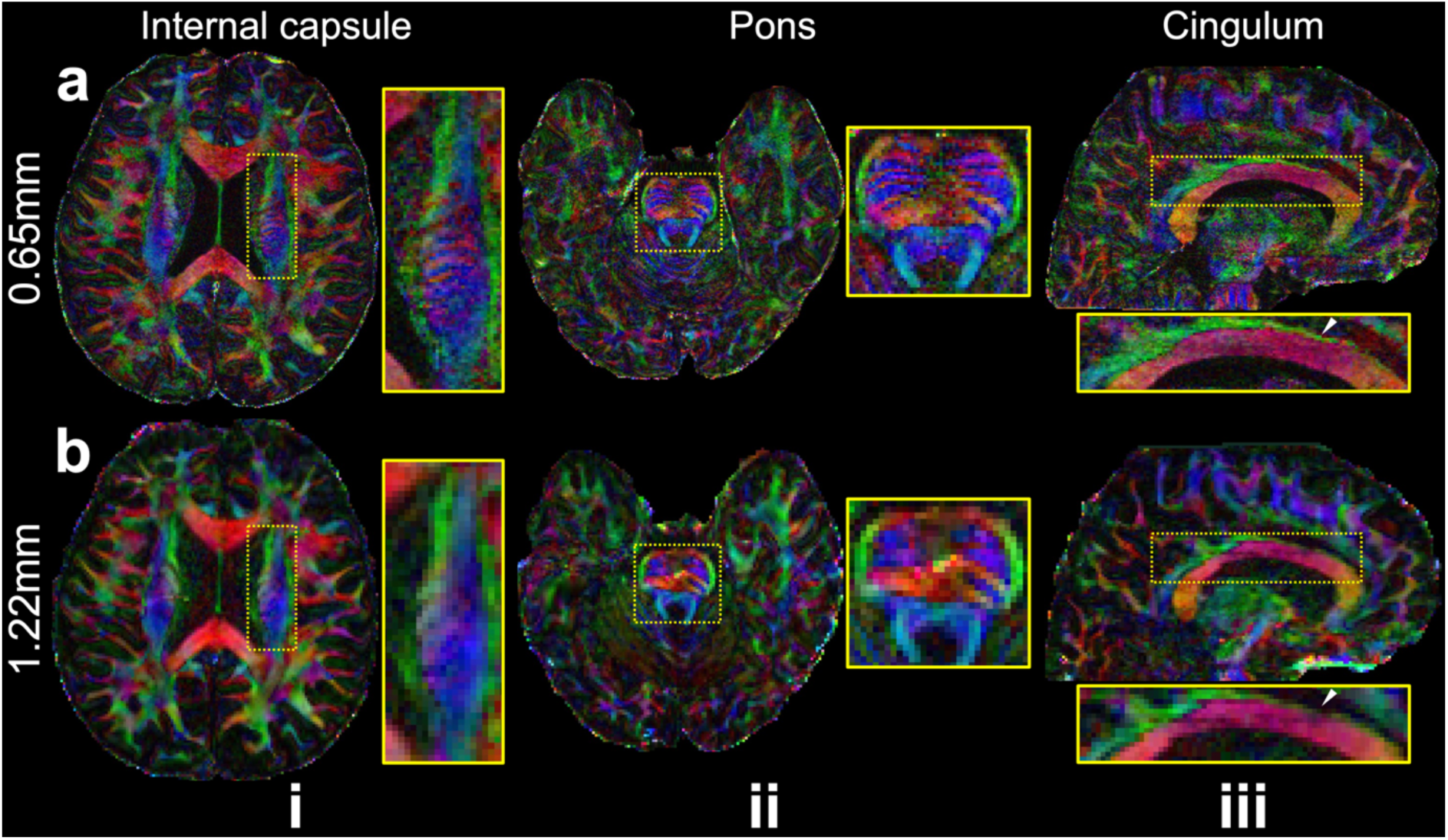
Comparison of 3T 0.65 mm and 1.22 mm isotropic resolution DTI. Two axial slices showing the internal capsule (i) and pons (ii), and a sagittal slice showing the cingulum bundle (iii) of FA modulated V1 of 0.65mm (a) and 1.22mm (b) isotropic resolution in-vivo diffusion data acquired at 3T are demonstrated. White arrows highlight the improved delineation of the cingulum bundle in the 0.65mm data.

Figure 6 shows the whole-brain diffusion data from 3T 0.53 mm Protocol. Across both subjects, our proposed acquisition and reconstruction framework consistently achieves high image quality with adequate SNR (Fig. 6a, c) and superior anatomical fidelity (Fig. 6b) at 0.53 mm isotropic resolution.

**Figure 6.**
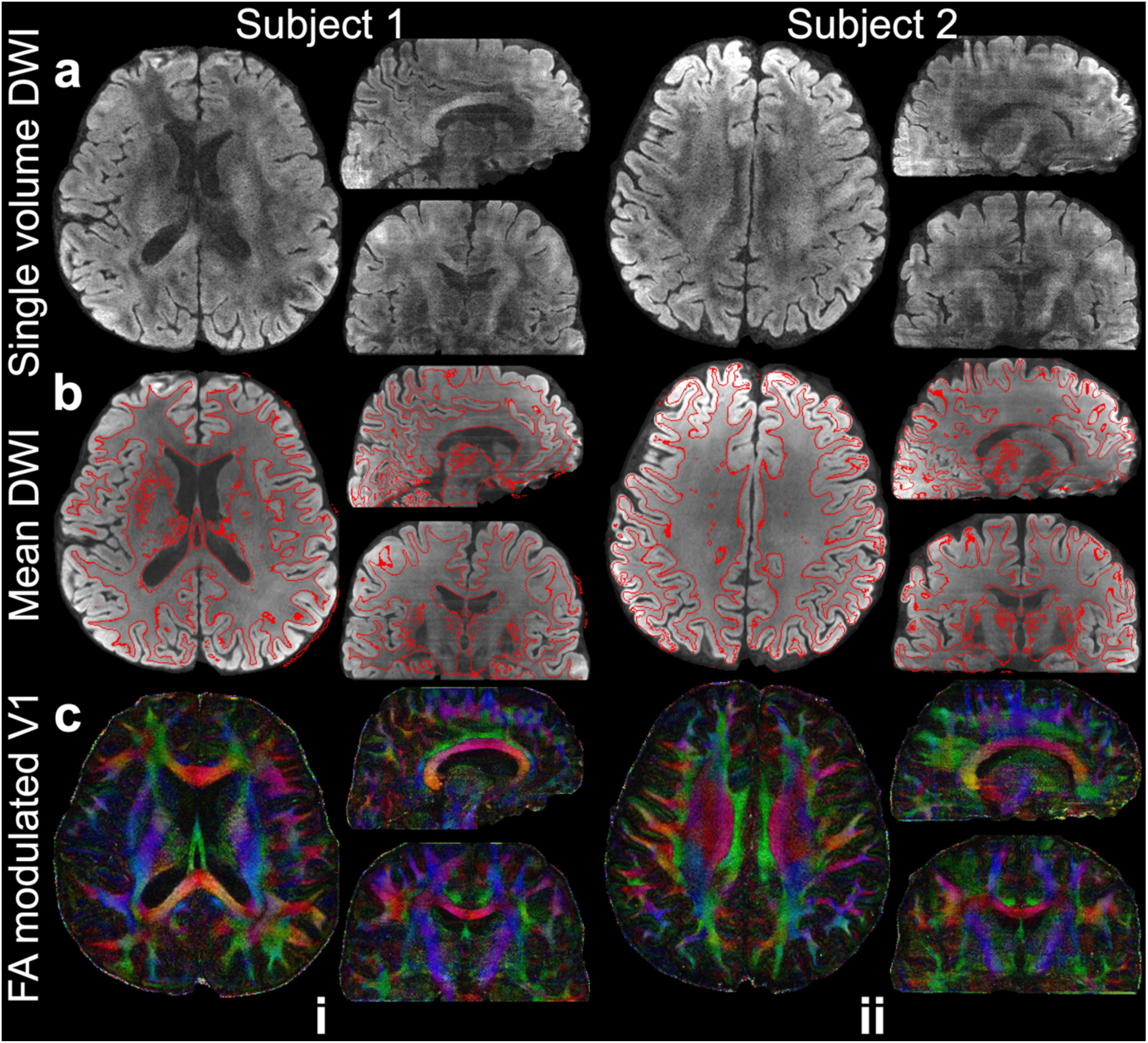
Whole-brain data from 3T 0.53 mm Protocol. Whole-brain in-vivo diffusion data of two subjects (b=1000 s/mm^2^) including diffusion-weighted image (DWI) along (0.5, -0.86, 0.02) (a), 20-direction mean DWI with gray-white matter boundary derived from the T1w image using FSL’s “fast” overlayed (b), FA modulated V1 (c) at 0.53 mm isotropic resolution are shown.

The superior quality of the 0.53 mm data is further demonstrated by the successful reconstruction of three representative white matter tracts (Fig. 7). Specifically, our data enable the tracking of the acoustic radiation, which crosses several major brain fiber systems with complex fiber crossings, making it difficult to reconstruct from in-vivo data (Maffei et al., 2018). The tracking results for the acoustic radiation on the 0.53 mm data are consistent with those from the 1.22 mm data (Fig. 7, i). The cingulate gyrus portion of the cingulum, which predominantly follows an anterior-posterior trajectory, is also successfully reconstructed in both datasets. The tract appears more extensive in the retrosplenial region in the 0.53 mm data, likely due to the higher resolution (Fig. 7, ii, yellow arrows). For the corticospinal tract, which primarily runs along the slice selection direction, the 0.53 mm data reveal clearer branching (Fig. 7, iii, yellow arrows at the top) and a thicker appearance near the pons (Fig. 7, iii, yellow arrows near the bottom). The clearer branching is likely due to the improved delineation of crossing fibers thanks to the higher resolution. The thicker tract near the pons is attributed to reduced distortion and better anatomical fidelity in the 0.53 mm data compared to the 1.22 mm data. Notably, the tractography utilized masks from FSL’s “autoPtx”, which is designed for conventional resolution fiber tracking (1-2 mm). It is expected that the advantage of high-resolution data would be even more pronounced with tailored tractography methods designed for such spatial resolution.

**Figure 7.**
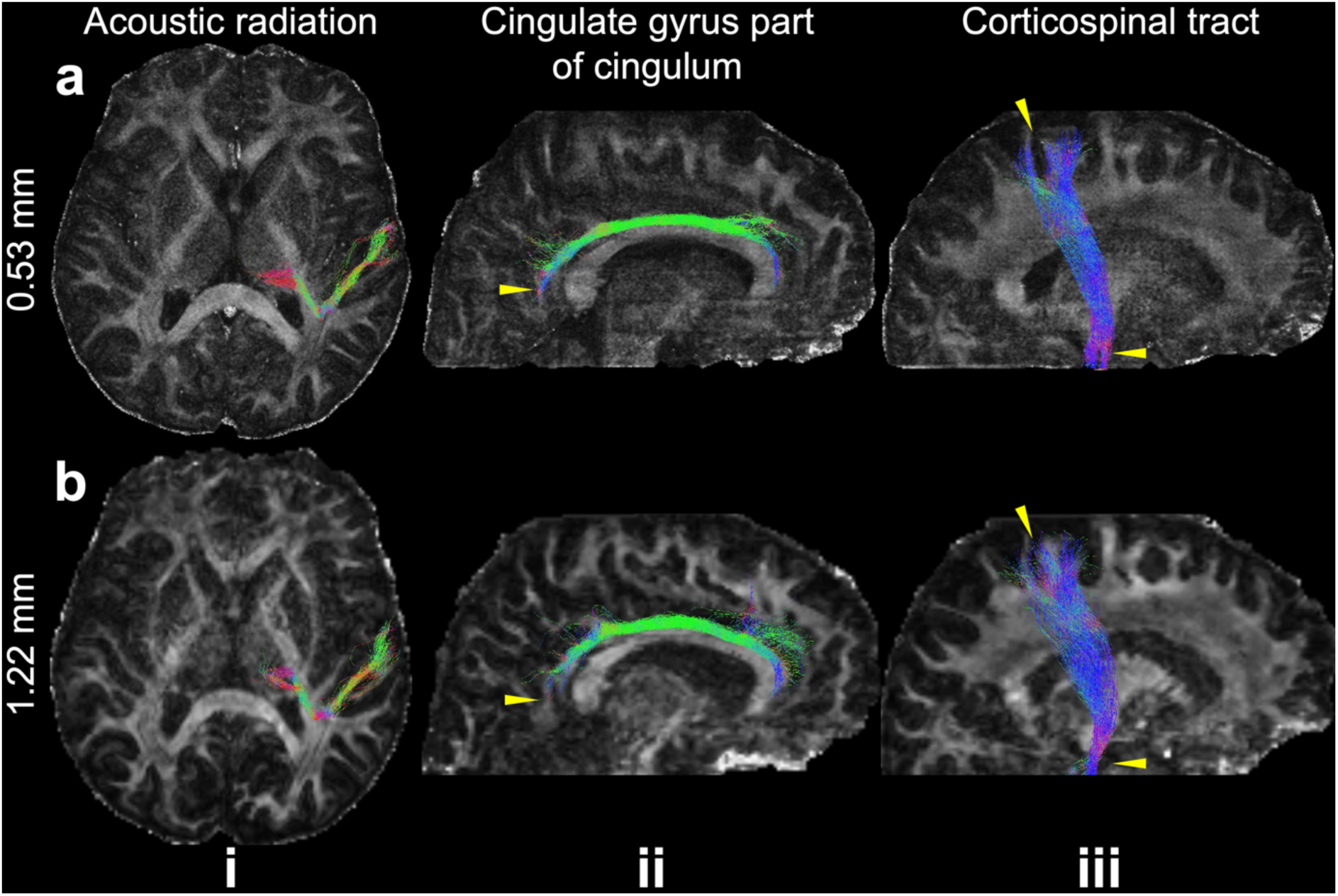
Example white matter tracts from the 0.53 mm and 1.22 mm data. The maximum intensity projections of three representative white matter tracts including acoustic radiation (i), cingulate gyrus part of cingulum (ii), and corticospinal tract (iii) are overlayed on the fractional anisotropy maps of 0.53 mm (a) and 1.22 mm (b) datasets. The yellow arrows indicate the region where the tractography on 0.53 mm data shows improvement compared to that on 1.22 mm data.

The 0.53 mm data also effectively reduce the gyral bias problem, particularly in small gyri (Fig. 8). In the 1.22 mm data, gyral bias is evident, with streamlines often terminating at the gyral crowns (Fig. 8, ii, iv). In contrast, the 0.53 mm data show the expected fanning pattern, with a greater number of streamlines extending to the gyral walls (Fig. 8, i, iii). This more even distribution of streamline termination points between gyral crowns and walls in the 0.53 mm data indicates a significant reduction in gyral bias with higher spatial resolution, consistent with previous post-mortem findings (Tendler et al., 2022). However, we found the gyral bias problem in our data is less pronounced in larger gyri, where the expected fanning pattern is also visible in the 1.22 mm data (Supplementary Fig. 4).

**Figure 8.**
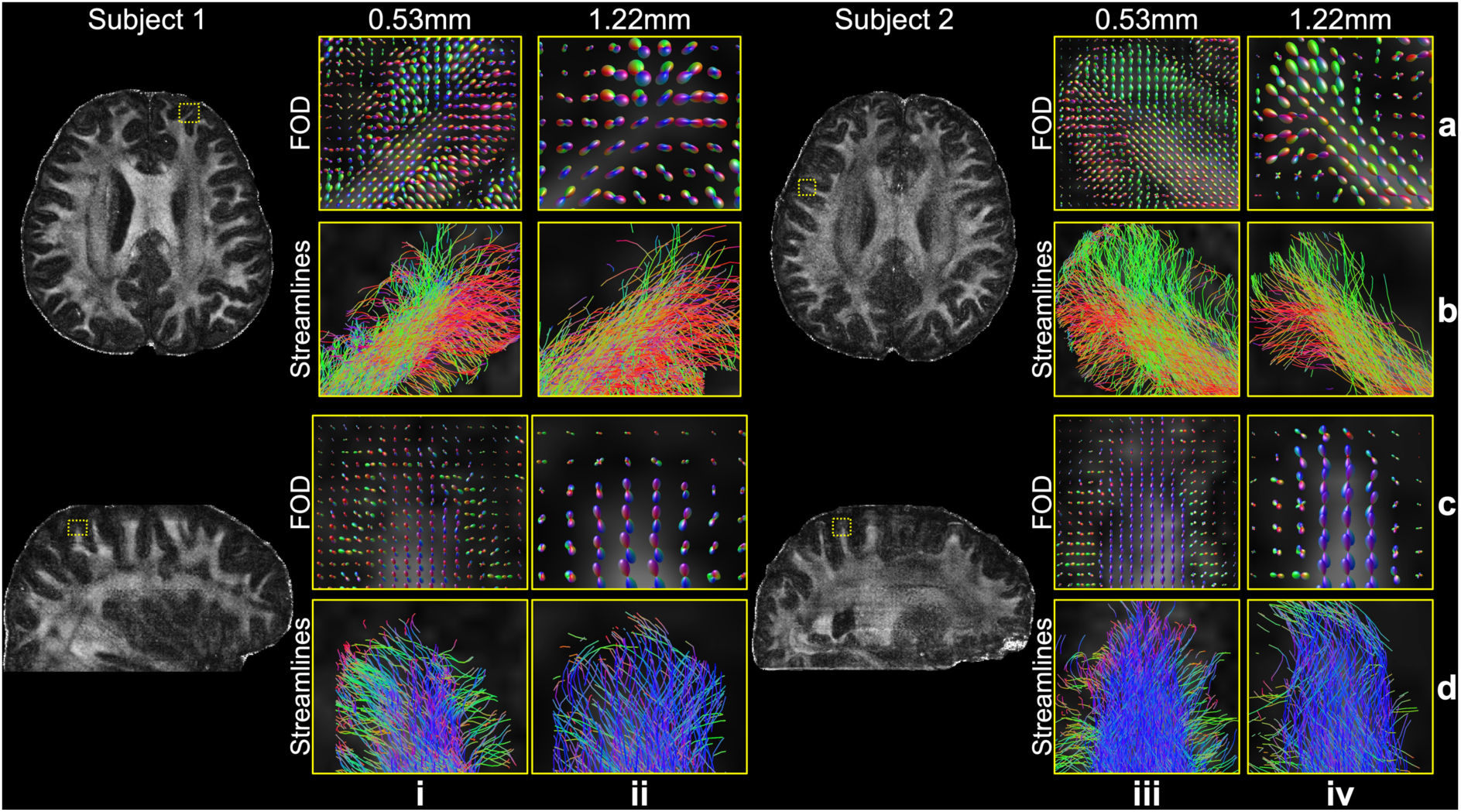
Gyral bias comparison between 0.53 mm and 1.22 mm data. The fiber orientation distributions (FOD) (a, c) and tractography streamlines (b, d) for representative gyri from the 0.53 mm (i, iii) and 1.22 mm (ii, iv) data of two subjects are shown to demonstrate the reduced gyral bias on high-resolution data. The fractional anisotropy maps at 0.53 mm resolution are also displayed to indicate the anatomical position of these gyri.

Compared to the 1.22 mm data, the 0.53 mm data also provide a more accurate mapping of U-fibers (Fig. 9). Our tractography setup successfully identified short association fibers in both datasets (Fig. 9a, b). When using one seed per voxel for both, the 0.53 mm data reveal denser whole-brain short association fibers, likely due to the substantially increased voxel number. The distribution of short association fibers in the 0.53 mm data is also more continuous along the gray-white matter boundary. These findings are consistent with previous studies (Song et al., 2014). Remarkably, the 0.53 mm data is able to resolve U-fibers even at sharp turnings along the gray-white matter boundary (Fig. 9c-e, i, ii), whereas the 1.22 mm data struggle to capture these fibers (Fig. 9c-e, iii, iv), even with increased seed numbers to compensate the voxel number difference (Fig. 9c-e, v). The tractography on 0.53 mm data with one seed per voxel and that on 1.22 mm data with 12 seeds per voxel produced comparable numbers of streamlines (1.75M vs. 1.84M). This confirms the improvement on the 0.53 mm data originates from the better resolved fiber turning curvature due to the higher spatial resolution, rather than simply an increase in seed or streamline numbers.

**Figure 9.**
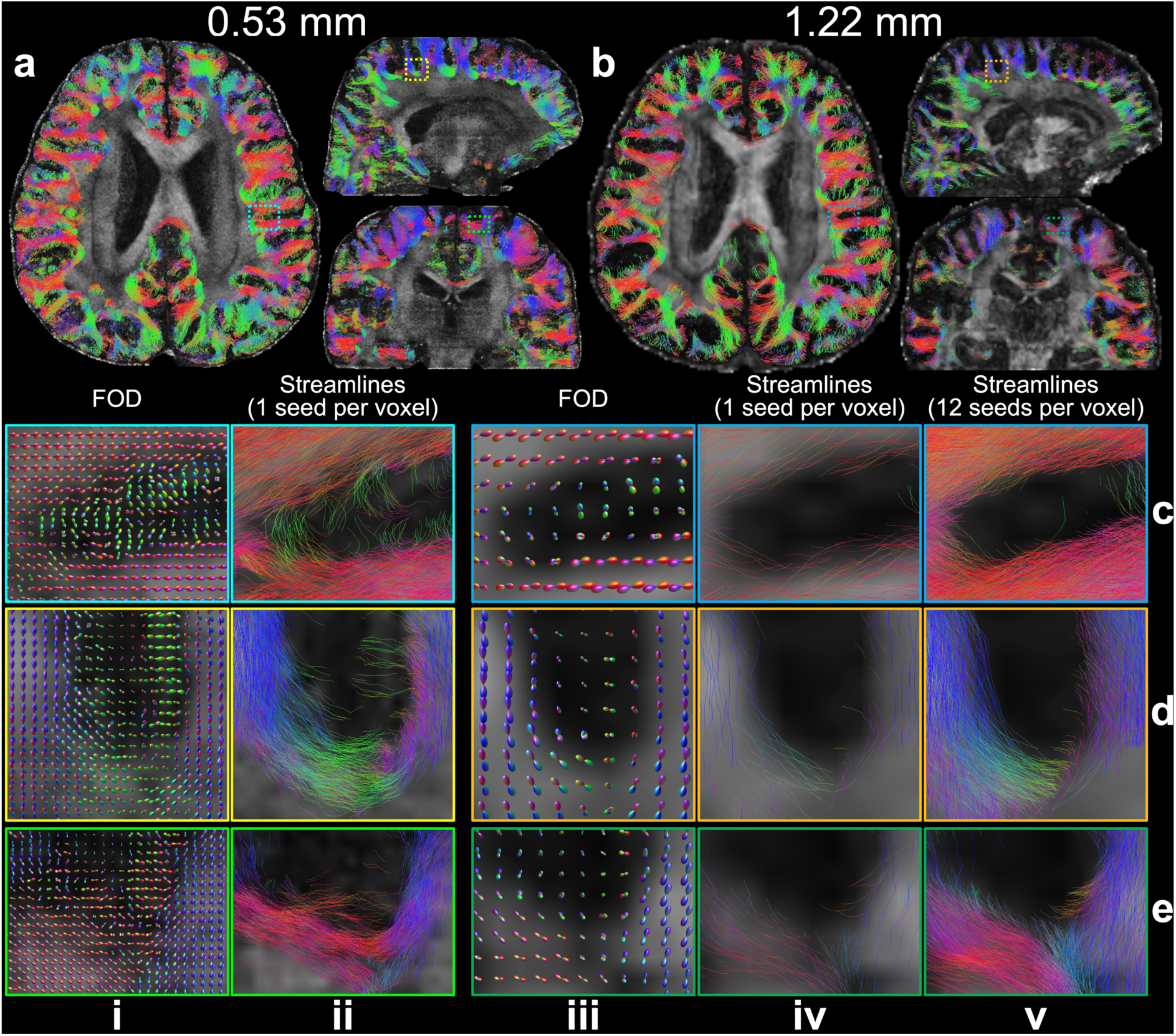
U-fibers comparison between 0.53 mm and 1.22 mm data. The whole-brain short association fibers tracked with one seed per voxel for 0.53 mm (a) and 1.22 mm (b) data are overlayed on fractional anisotropic maps. Representative enlarged regions (c-e) show the fiber orientation distributions (FOD) (i, iii), the tracked streamlines with one seed per voxel (ii, iv) for the 0.53 mm (i, ii) and 1.22 mm (iii, iv) data, and the tracked streamlines with 12 seeds per voxel for the 1.22 mm data (v) to compensate the difference in voxel numbers due to resolution.

Our acquisition and reconstruction framework shows robustness at higher field strength of 7T (Fig. 10). The diffusion data from the 7T 0.61 mm Protocol exhibit excellent SNR even with only six diffusion encoding directions, thanks to the SNR benefit at 7T (Supplementary Fig. 3). Compared to the 1.05 mm isotropic resolution data, the submillimeter dataset reveals significantly more detailed microstructure, including better resolution of fanning patterns in the gyri and clearer delineation of small structures (Fig. 10, white arrows). Notably, our in-vivo results for resolving transverse pontine fibers are in excellent agreement with previous post-mortem data acquired with much longer scan times on the same 7T scanner (Tendler et al., 2022).

**Figure 10.**
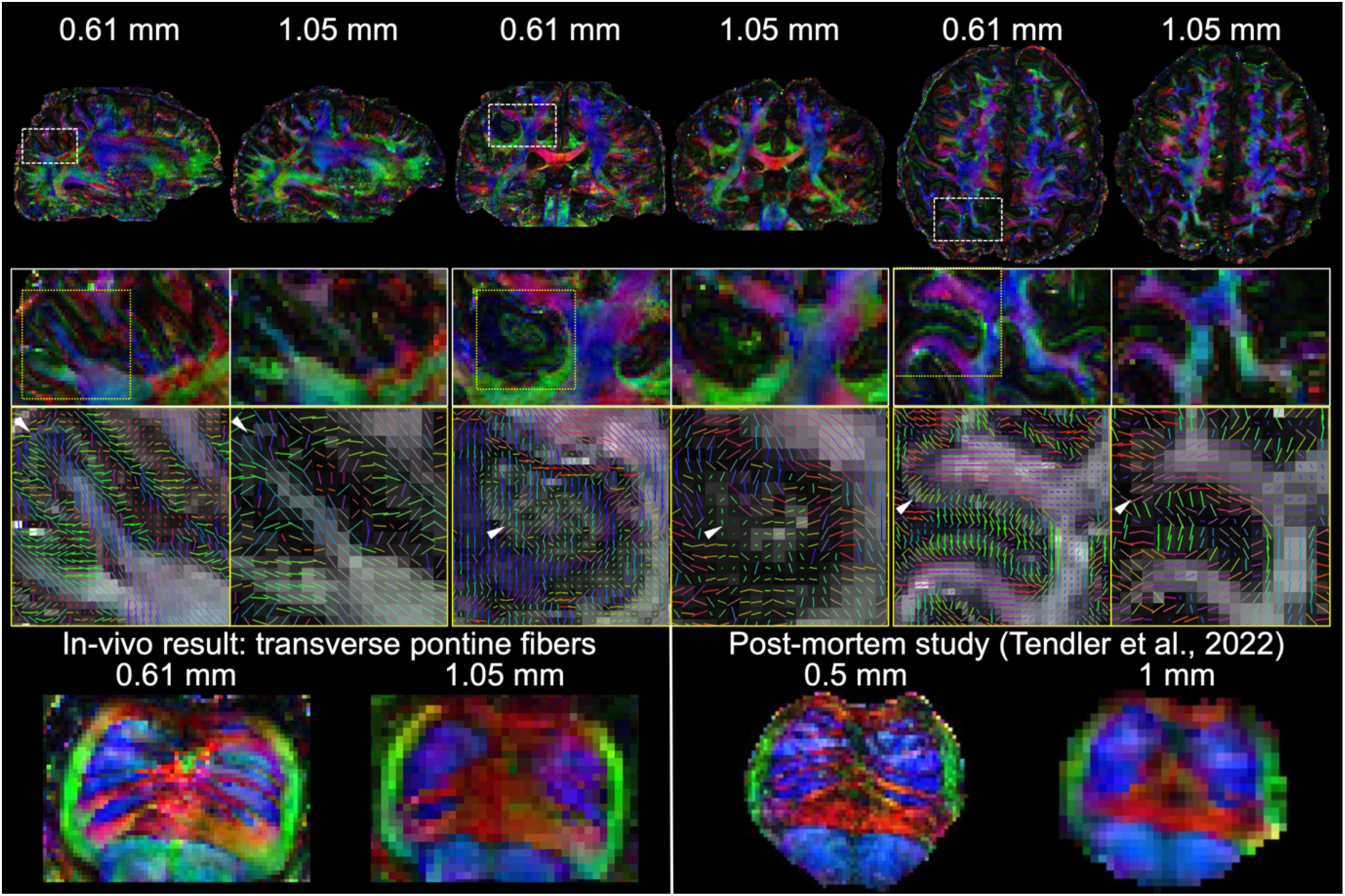
Comparisons of 7T 0.61 mm and 1.05 mm diffusion data. The sagittal, coronal, and axial slices of fractional anisotropy (FA) modulated V1 and their enlarged regions of 0.61 mm and 1.05 mm isotropic resolution in-vivo diffusion data acquired at 7T are demonstrated, with white arrows highlighting the more detailed microstructure resolved by the high resolution at 0.61 mm. The traverse pontine fibers demonstrate similar patterns compared to previous post-mortem study (Tendler et al., 2022) at 0.5 mm isotropic resolution from the same scanner.

## 4. Discussion

This study introduces an acquisition and reconstruction framework to achieve high-quality submillimeter dMRI. Our acquisition takes advantage of the optimal SNR efficiency provided by 3D multi-slab EPI, with in-plane segmentations integrated to shorten the effective echo spacing, readout, and TE for reduced distortion, T2* blurring, and SNR penalty. The sampling order is designed for improved robustness against subject motion. A denoiser-regularized reconstruction approach is proposed to suppress noise while maintaining consistency with the acquired data to reduce bias or blurring. Our in-vivo experiments at 3T demonstrate that our method consistently produces high-quality diffusion data at 0.65 mm and 0.53 mm isotropic resolutions. The analyses of these submillimeter datasets reveal enhanced microstructural detail, mitigating the “gyral bias” problem and improving the precision of U-fiber tracking compared to data acquired at conventional resolutions of 1.22 mm from the same subjects at the same scanner. The framework also proves effective at 7T, delivering similarly robust results. These findings suggest that the proposed framework represents a promising advancement in high-resolution dMRI, offering the potential for more accurate characterization of brain microstructure and expanded applications in neuroscientific research.

The optimization of our sampling strategies considers the effective resolution, SNR, distortion, scan time, and motion robustness. Compared to previous studies that achieve submillimeter resolutions using super-resolution methods (Dong et al., 2024; Liao et al., 2023; Setsompop et al., 2018), our method using 3D multi-slab EPI has several key advantages: our method using 3D multi-slab EPI offers several key advantages: (i) superior SNR efficiency enabled by short TR and orthogonal encoding bases from Fourier encoding, and (ii) high-fidelity voxel shapes produced directly by Fourier encoding, avoiding the need for Tikhonov regularization used in super-resolution reconstruction. However, for EPI-based sampling, higher resolution imaging requires longer echo spacing, readout, and TE, and therefore more severe distortion, T2* blurring, and T2 signal decay. For example, for *N*_*seg*_=3 sampling without PF at 3T, image displacement (assuming a 50 Hz B0 offset), image blurring ((effective resolution-nominal resolution)/nominal resolution) and T2 signal decay (*e*^−*TE*/*T*2^) for 0.6 mm imaging are 4.4 mm, 41.3%, and 0.103, respectively, compared to 3.2 mm, 25.3%, and 0.293 for 1 mm imaging (Fig. 1). This requires a larger *N*_*seg*_ for submillimeter imaging to improve effective resolution, SNR, and minimize distortion, though at the cost of longer scan times. In the 3T 0.65 mm Protocol, we acquired fully-sampled data with *N*_*seg*_=6 without using PF, which achieved superior SNR, image sharpness, and anatomical fidelity (Fig. 4). To enable more advanced diffusion analyses with more diffusion encoding directions, we explored the potential for acceleration by acquiring fewer segments (Fig. 3) and achieved *R*_*eff*_=2 in the 3T 0.53 mm Protocol without significantly compromising the image quality, allowing for the acquisition of 20 diffusion encoding directions in a one-hour session. Notably, the TR of 2s in this protocol is within the SNR-efficient range. Additionally, we applied a motion-robust sampling order for 3D EPI (Supplementary Fig. 1), which particularly improves the b=0 images where motion artifacts are more noticeable due to strong CSF signal aliasing.

Our denoiser-regularized reconstruction DnSPIRiT effectively suppresses noise while maintaining image fidelity by enforcing consistency with the acquired data. Essentially, the denoised image serves as a regularization to suppress noise during reconstruction, instead of directly being used as the final output. The experiment shows our method effectively reduces the noise without introducing significant blurring or bias (Fig. 2), supported by the fact that the removed noise generally exhibits random distribution without noticeable anatomical structures (Fig. 2e). DnSPIRiT proves even more beneficial in under-sampled reconstruction where the noise level is higher (Fig. 3). The plug-and-play nature of DnSPIRiT allows for the integration of advanced denoisers without adding significant computational burden in directly solving Eq. 2, which is crucial for high-resolution dMRI reconstruction due to the large data size. We demonstrated its compatibility with BM4D for 6-direction data denoising (3T 0.65 mm Protocol, 7T 0.61 mm Protocol) and NORDIC for 20-direction data denoising (3T 0.53 mm Protocol). Incorporating recently developed deep learning-based dMRI denoisers (Fadnavis et al., 2020; Tian et al., 2020; Tian et al., 2022) that outperform conventional methods is expected to further improve the reconstruction.

The comprehensive in-vivo experiments comparing submillimeter data with prospectively acquired conventional-resolution data (1.22 mm and 1.05 mm) validate the benefits of higher spatial resolution for diffusion analyses. In previous studies (Chang et al., 2015; Song et al., 2014; Wang et al., 2021), the comparisons between the high-resolution and low-resolution data were based on retrospective under-sampling. One limitation of those comparisons is that the increased image blurring and SNR penalty associated with high-resolution imaging (Fig. 1, Supplementary Fig. 3) can propagate into the under-sampled low-resolution data, leading to lower image quality than what can be practically achieved. Instead, we prospectively acquired data at 1.22 mm and 1.05 mm with practical imaging protocols from the same scanners as the submillimeter resolution data. Using such approach, the datasets at conventional resolutions also demonstrate excellent data quality. For instance, the expected fiber fanning pattern can be resolved at large gyri from the 1.22 mm data (Supplementary Fig. 4). These prospective acquisitions provide a more realistic comparison, revealing that submillimeter data still significantly outperform 1.22 mm data in reducing gyral bias at small gyri and mapping U-fibers in sharp gray-white matter boundary turnings. Moreover, our results’ strong agreement with previous post-mortem findings (Tendler et al., 2022) on the same 7T scanner (Fig. 10) further reinforces the reliability of these comparisons.

Our implementation of the sequence using Pulseq enhances the accessibility of our method. The scanner-agnostic, open-source nature of Pulseq facilitates the straightforward translation of our framework to different scanner platforms. Demonstrating the ability to produce high-quality data on a clinical scanner (Siemens 3T Prisma) highlights the potential for broader application of our methods across various clinical and research settings. We further demonstrated the method’s compatibility a higher field strength at 7T (Siemens 7T Magnetom), leveraging the SNR advantage (Fig. 10). Additionally, the state-of-the-art high-performance scanners with stronger gradients, such as the Siemens 3T Connectome 2.0 (Huang et al., 2021), GE 3T Ultra-High Performance, and Siemens 7T NexGen (Feinberg et al., 2023), offer promising opportunities to further reduce readout times and TE, improving image quality and potentially enabling even higher spatial resolutions.

Robust acquisition of submillimeter dMRI data opens up new possibilities for medical image analysis and neuroscientific research. Our study demonstrates its potential to address “gyrus bias” and improve U-fiber mapping, leading to more accurate reconstruction of intra-and inter-gyri connections. Future work could leverage the rich detail of ultrahigh-resolution data through more quantitative analyses. One potential direction is integrating cortical curvature from T1w images to enable precise whole-brain gyri segmentation to improve whole-brain U-fiber mapping at submillimeter resolution. Submillimeter dMRI may also aid in mapping small but crucial subcortical structures targeted for deep brain stimulation (Maffei et al., 2022; Saleem et al., 2021). Public subcortical atlases (e.g., https://www.lead-dbs.org/helpsupport/knowledge-base/atlasesresources/atlases-2/) offer useful resources for defining seeding and targeting masks for tractography to map white matter circuits in these regions.

One limitation of our method is the relatively long scan and reconstruction times. The scan time per volume (2.7 minutes in the 3T 0.53 mm Protocol) restricts the number of diffusion encoding directions. Further acceleration could be achieved by leveraging k-q space joint reconstruction, which shares information across diffusion directions (Wu et al., 2019). Additionally, recent advances in eliminating navigator acquisition in 3D multi-slab EPI (Li et al., 2024) could be incorporated into our framework to further enhance scan efficiency. On the reconstruction side, processing submillimeter data remains time-intensive due to the large matrix size (∼12 hours per slab on a single CPU for data from the 3T 0.53 mm Protocol). Model-based deep learning reconstruction methods (Aggarwal et al., 2018) present a promising avenue to significantly reduce computational time and potentially further improve SNR.

## 5. Conclusion

In this work, a novel acquisition and reconstruction framework is proposed to achieve high-quality submillimeter dMRI 0.5-0.6 mm isotropic resolution for in-vivo human brains. Comprehensive in-vivo experiments demonstrate the superior SNR, image sharpness, and anatomical fidelity of the submillimeter data, which reveal substantially more microstructure details, reduce the gyral bias, and improve the U-fiber mapping compared to data at conventional resolutions (1-1.2 mm). The method also shows robustness at 7T, where data show excellent agreement with previous post-mortem data at 0.5 mm resolution acquired from the same scanner. With the scanner-agnostic, open-source implementation using Pulseq, our approach holds promise in benefitting a wide range of medical image analyses and neuroscientific research.

## Acknowledgement

The authors would like to thank Profs. Qiyuan Tian and Hua Guo at Tsinghua University for the helpful discussions on high-resolution diffusion data analysis, and Dr. Fredrick Lange at the University of Oxford for his insights on running FSL’s “eddy” on high-resolution diffusion data. We are also deeply grateful to the Pulseq and Harmonized MRI teams, led by Profs. Maxim Zaitsev, Jon-Fredrik Nielsen, Yogesh Rathi, and Berkin Bilgic, for their assistance with Pulseq programming and their dedication to the development, optimization, and open-source sharing of Pulseq. W.W. is supported by the Royal Academy of Engineering (RF\201819\18\92). K.L.M. is supported by the Wellcome Trust (WT202788/Z/16/A). This study is supported by the NIHR Oxford Health Biomedical Research Centre (NIHR203316). The views expressed are those of the author(s) and not necessarily those of the NIHR or the Department of Health and Social Care. The Wellcome Centre for Integrative Neuroimaging is supported by core funding from the Wellcome Trust (203139/Z/16/Z and 203139/A/16/Z). For the purpose of Open Access, the authors have applied a CC BY public copyright license to any Author Accepted Manuscript (AAM) version arising from this submission.

## Supplementary Information

**Supplementary Figure 1.**
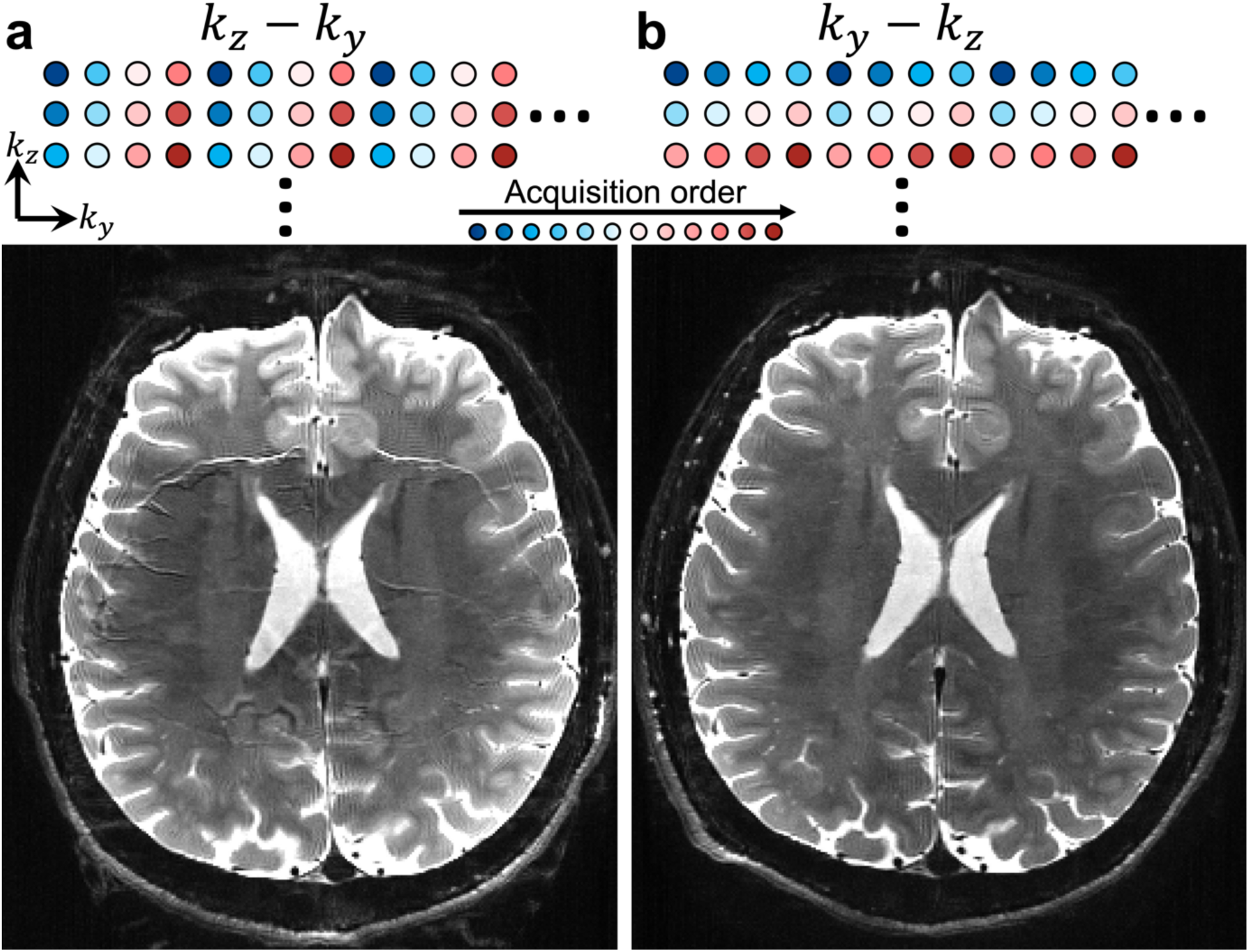
Impact of the sampling order of in-plane segmented 3D EPI. The “kz-ky” (a, all kz planes for one ky segment are acquired before proceeding to the next segment) and “ky-kz” (b, all ky segments for one kz plane are acquired before proceeding to the next kz) sampling orders are illustrated with a simplified 4-segment example. Representative b=0 images (0.65 mm isotropic resolution, 6-segment acquisition) are demonstrated below the corresponding sampling orders.

**Supplementary Figure 2.**
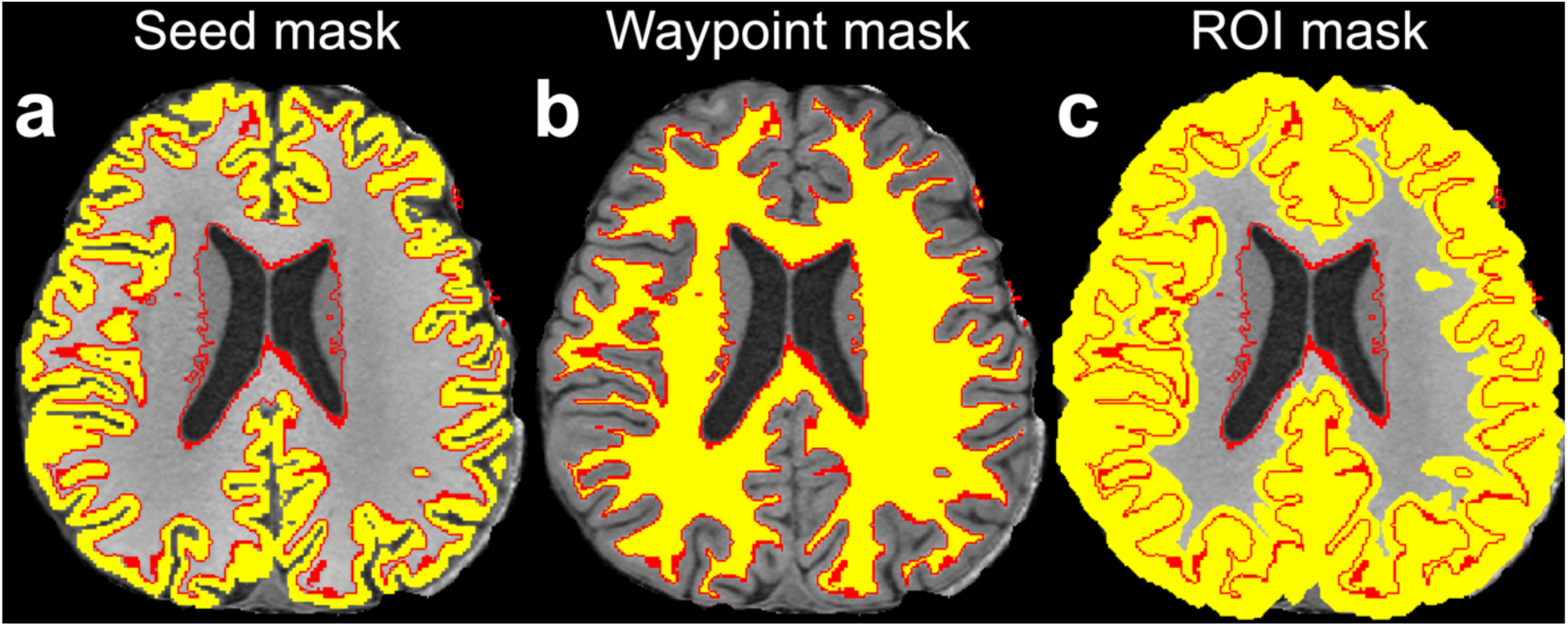
Masks for tracking the short association fibers. The seed (a), waypoint (b), and region-of-interest (ROI) (c) masks (yellow) for tracking the short association fibers are displayed on an axial slice of the T1w image, with the white matter boundary marked in red and overlayed.

**Supplementary Figure 3.**
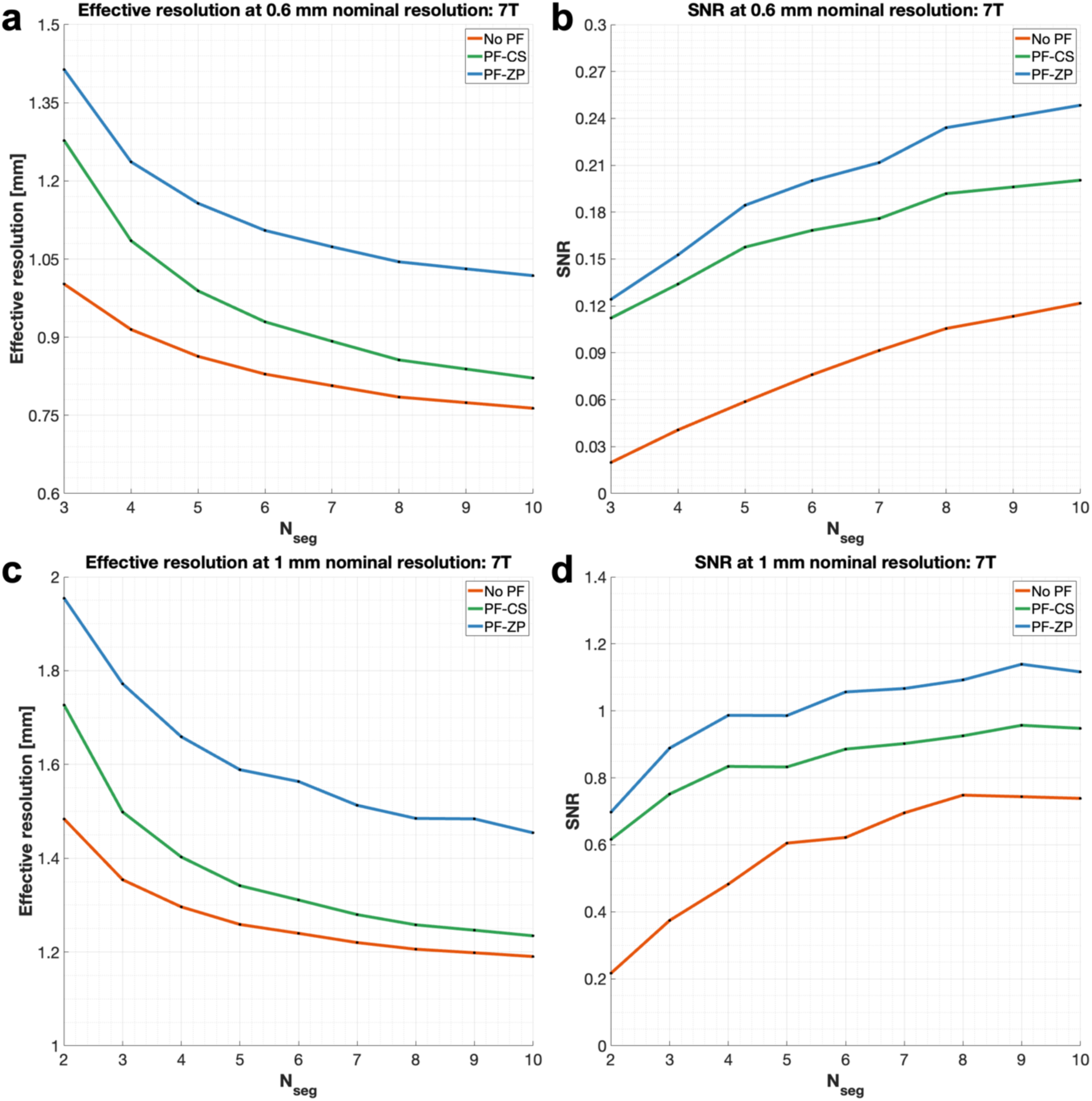
Simulation of effective resolution and SNR for high-resolution diffusion-weighted EPI at 7T. No partial Fourier (No PF, red), 6/8 PF with conjugate symmetric filling (PF-CS, green), and 6/8 PF with zero padding (PF-ZP, blue) samplings are investigated at 1 mm (a, b) and 0.6 mm (c, d) for white matter at 7T (TR=3s, b-value=1000 s/mm^2^, bandwidths = 992/1384 Hz/pixel for 0.6/1 mm acquisitions, T1/T2/T2*=1200/47/26.8 ms) for different in-plane segmentation numbers (*N*_*seg*_).

**Supplementary Figure 4.**
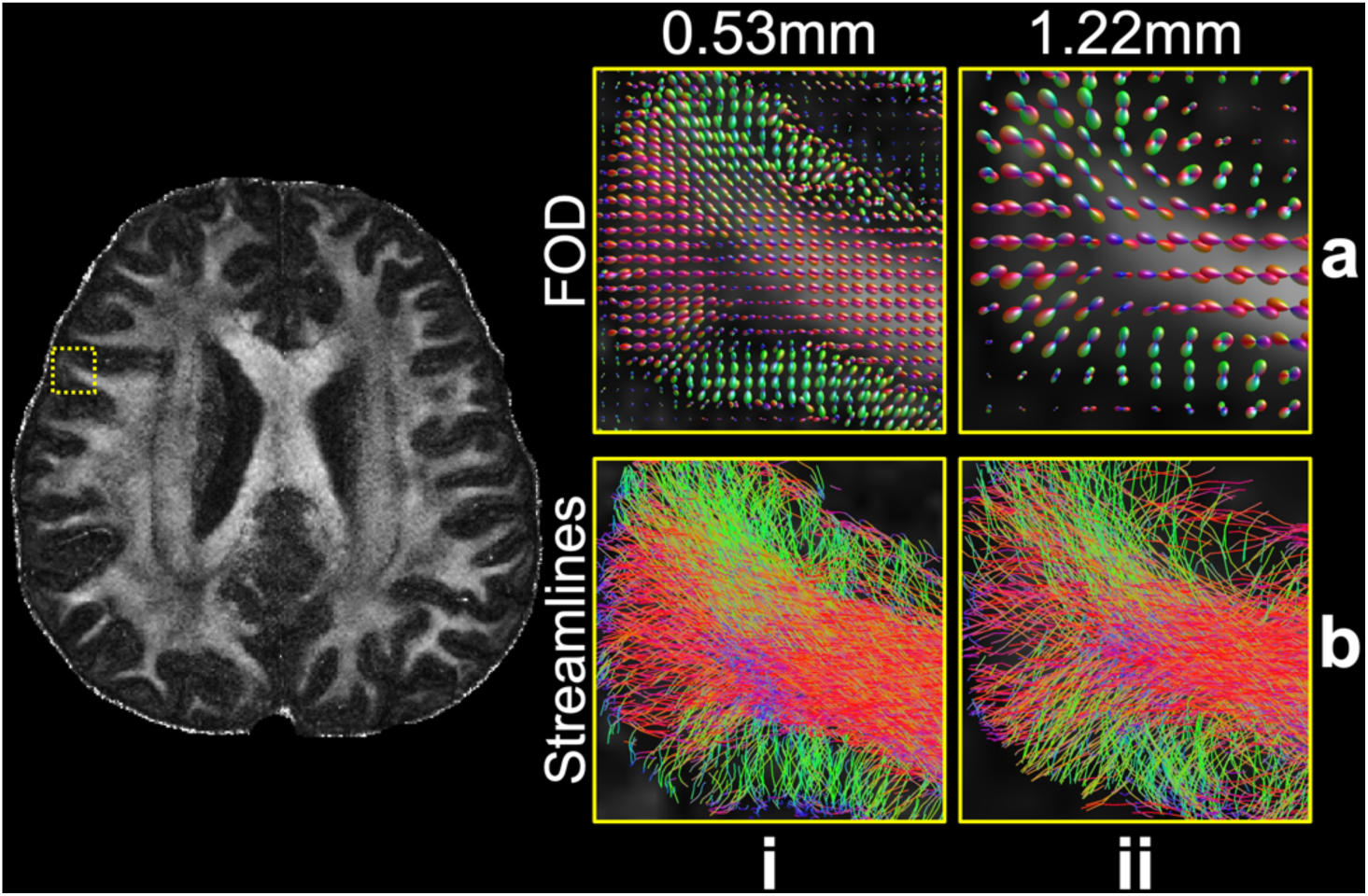
Fiber distributions at a large gyrus. The fiber orientation distributions (FOD) (a) and tractography streamlines (b) for a large gyrus from the 0.53 mm (i) and 1.22 mm (ii) data are shown. At this large gyrus, the gyral bias problem is less pronounced even on 1.22 mm data. The fractional anisotropy (FA) map at 0.53 mm resolution is also displayed to indicate the anatomical position of the gyrus.

## Notes

### Competing Interest Statement

The authors have declared no competing interest.

## Reference

Aggarwal, H. K., Mani, M. P., & Jacob, M. (2018). MoDL: Model-based deep learning architecture for inverse problems. IEEE transactions on medical imaging, 38(2), 394–405.

Ahmad, R., Bouman, C. A., Buzzard, G. T., Chan, S., Liu, S., Reehorst, E. T., & Schniter, P. (2020). Plug-and-play methods for magnetic resonance imaging: Using denoisers for image recovery. IEEE signal processing magazine, 37(1), 105–116.

Andersson, J. L., Skare, S., & Ashburner, J. (2003). How to correct susceptibility distortions in spin-echo echo-planar images: application to diffusion tensor imaging. NeuroImage, 20(2), 870–888.

Andersson, J. L., & Sotiropoulos, S. N. (2016). An integrated approach to correction for off-resonance effects and subject movement in diffusion MR imaging. NeuroImage, 125, 1063–1078.

Bautista, T., O’Muircheartaigh, J., Hajnal, J., & J-Donald, T. (2021). Removal of Gibbs ringing artefacts for 3D acquisitions using subvoxel shifts. Proc Int Soc Magn Reson Med, Virtual Conference.

Behrens, T. E., Berg, H. J., Jbabdi, S., Rushworth, M. F., & Woolrich, M. W. (2007). Probabilistic diffusion tractography with multiple fibre orientations: What can we gain? NeuroImage, 34(1), 144–155.

Behrens, T. E., Woolrich, M. W., Jenkinson, M., Johansen-Berg, H., Nunes, R. G., Clare, S., Matthews, P. M., Brady, J. M., & Smith, S. M. (2003). Characterization and propagation of uncertainty in diffusion-weighted MR imaging. Magnetic Resonance in Medicine: An Official Journal of the International Society for Magnetic Resonance in Medicine, 50(5), 1077–1088.

Buades, A., Coll, B., & Morel, J.-M. (2011). Non-local means denoising. Image Processing On Line, 1, 208–212.

Chang, H.-C., Sundman, M., Petit, L., Guhaniyogi, S., Chu, M.-L., Petty, C., Song, A. W., & Chen, N.-k. (2015). Human brain diffusion tensor imaging at submillimeter isotropic resolution on a 3 Tesla clinical MRI scanner. NeuroImage, 118, 667–675.

Chen, N.-k., Guidon, A., Chang, H.-C., & Song, A. W. (2013). A robust multi-shot scan strategy for high-resolution diffusion weighted MRI enabled by multiplexed sensitivity-encoding (MUSE). NeuroImage, 72, 41–47.

Cottaar, M., Bastiani, M., Boddu, N., Glasser, M. F., Haber, S., Van Essen, D. C., Sotiropoulos, S. N., & Jbabdi, S. (2021). Modelling white matter in gyral blades as a continuous vector field. NeuroImage, 227, 117693.

Dabov, K., Foi, A., Katkovnik, V., & Egiazarian, K. (2007). Image denoising by sparse 3-D transform-domain collaborative filtering. IEEE Transactions on Image Processing, 16(8), 2080–2095.

Dai, E., Liu, S., & Guo, H. (2021). High-resolution whole-brain diffusion MRI at 3T using simultaneous multi-slab (SMSlab) acquisition. NeuroImage, 237, 118099.

Dong, Z., Reese, T. G., Lee, H.-H., Huang, S. Y., Polimeni, J. R., Wald, L. L., & Wang, F. (2024). Romer-EPTI: Rotating-view motion-robust super-resolution EPTI for SNR-efficient distortion-free in-vivo mesoscale dMRI and microstructure imaging. bioRxiv.

Engström, M., & Skare, S. (2013). Diffusion-weighted 3D multislab echo planar imaging for high signal -to - noise ratio efficiency and isotropic image resolution. Magnetic resonance in medicine, 70(6), 1507–1514.

Fadnavis, S., Batson, J., & Garyfallidis, E. (2020). Patch2Self: Denoising Diffusion MRI with Self-Supervised Learning. Advances in Neural Information Processing Systems, 33, 16293–16303.

Feinberg, D. A., Beckett, A. J., Vu, A. T., Stockmann, J., Huber, L., Ma, S., Ahn, S., Setsompop, K., Cao, X., & Park, S. (2023). Next-generation MRI scanner designed for ultra-high-resolution human brain imaging at 7 Tesla. Nature methods, 20(12), 2048–2057.

Feizollah, S., & Tardif, C. L. (2023). High-resolution diffusion-weighted imaging at 7 Tesla: single-shot readout trajectories and their impact on signal-to-noise ratio, spatial resolution and accuracy. NeuroImage, 274, 120159.

Fischl, B. (2012). FreeSurfer. NeuroImage, 62(2), 774–781.

Frost, R., Miller, K. L., Tijssen, R. H., Porter, D. A., & Jezzard, P. (2014). 3D Multi-slab diffusion-weighted readout-segmented EPI with real-time cardiac-reordered k-space acquisition. Magnetic resonance in medicine, 72(6), 1565–1579.

Glasser, M. F., Smith, S. M., Marcus, D. S., Andersson, J. L., Auerbach, E. J., Behrens, T. E., Coalson, T. S., Harms, M. P., Jenkinson, M., & Moeller, S. (2016). The human connectome project’s neuroimaging approach. Nature neuroscience, 19(9), 1175–1187.

Griswold, M. A., Jakob, P. M., Heidemann, R. M., Nittka, M., Jellus, V., Wang, J., Kiefer, B., & Haase, A. (2002). Generalized autocalibrating partially parallel acquisitions (GRAPPA). Magnetic Resonance in Medicine: An Official Journal of the International Society for Magnetic Resonance in Medicine, 47(6), 1202–1210.

Haldar, J. P., & Zhuo, J. (2016). P - LORAKS: low - rank modeling of local k - space neighborhoods with parallel imaging data. Magnetic resonance in medicine, 75(4), 1499–1514.

Huang, S. Y., Witzel, T., Keil, B., Scholz, A., Davids, M., Dietz, P., Rummert, E., Ramb, R., Kirsch, J. E., & Yendiki, A. (2021). Connectome 2.0: Developing the next-generation ultra-high gradient strength human MRI scanner for bridging studies of the micro-, meso-and macro-connectome. NeuroImage, 243, 118530.

Ivanov, D., Barth, M., Uludağ, K., & Poser, B. A. (2015). Robust ACS acquisition for 3D echo planar imaging. Proceedings of the 23rd Annual Meeting of ISMRM, Toronto, Ontario, Canada.

Jenkinson, M., Bannister, P., Brady, M., & Smith, S. (2002). Improved optimization for the robust and accurate linear registration and motion correction of brain images. NeuroImage, 17(2), 825–841.

Jenkinson, M., Beckmann, C. F., Behrens, T. E., Woolrich, M. W., & Smith, S. M. (2012). Fsl. NeuroImage, 62(2), 782–790.

Jenkinson, M., & Smith, S. (2001). A global optimisation method for robust affine registration of brain images. Medical image analysis, 5(2), 143–156.

Layton, K. J., Kroboth, S., Jia, F., Littin, S., Yu, H., Leupold, J., Nielsen, J. F., Stöcker, T., & Zaitsev, M. (2017). Pulseq: a rapid and hardware - independent pulse sequence prototyping framework. Magnetic resonance in medicine, 77(4), 1544–1552.

Li, Z., Miller, K., Chen, X., Chiew, M., & Wu, W. (2024). Self-navigated 3D diffusion MRI using an optimized CAIPI sampling and structured low-rank reconstruction estimated navigator. IEEE transactions on medical imaging.

Li, Z., Miller, K. L., Andersson, J. L., Zhang, J., Liu, S., Guo, H., & Wu, W. (2023). Sampling strategies and integrated reconstruction for reducing distortion and boundary slice aliasing in high-resolution 3D diffusion MRI. Magnetic resonance in medicine, 90(4), 1484–1501.

Liao, C., Bilgic, B., Tian, Q., Stockmann, J. P., Cao, X., Fan, Q., Iyer, S. S., Wang, F., Ngamsombat, C., & Lo, W. C. (2021). Distortion-free, high-isotropic-resolution diffusion MRI with gSlider BUDA-EPI and multicoil dynamic B0 shimming. Magnetic resonance in medicine, 86(2), 791–803.

Liao, C., Yarach, U., Cao, X., Iyer, S. S., Wang, N., Kim, T. H., Tian, Q., Bilgic, B., Kerr, A. B., & Setsompop, K. (2023). High-fidelity mesoscale in-vivo diffusion MRI through gSlider-BUDA and circular EPI with S-LORAKS reconstruction. NeuroImage, 275, 120168.

Lustig, M., & Pauly, J. M. (2010). SPIRiT: iterative self - consistent parallel imaging reconstruction from arbitrary k-space. Magnetic resonance in medicine, 64(2), 457–471.

Maffei, C., Jovicich, J., De Benedictis, A., Corsini, F., Barbareschi, M., Chioffi, F., & Sarubbo, S. (2018). Topography of the human acoustic radiation as revealed by ex vivo fibers micro-dissection and in vivo diffusion-based tractography. Brain Structure and Function, 223, 449–459.

Maffei, C., Wang, F., Haber, S., & Yendiki, A. (2022). Submillimeter dMRI protocol optimization for accurate in-vivo reconstruction of deep-brain circuitry. Proceedings of the 30th Annual Meeting of ISMRM, London, UK.

Maggioni, M., Katkovnik, V., Egiazarian, K., & Foi, A. (2012). Nonlocal transform-domain filter for volumetric data denoising and reconstruction. IEEE Transactions on Image Processing, 22(1), 119–133.

Manzano Patron, J. P., Moeller, S., Andersson, J. L., Ugurbil, K., Yacoub, E., & Sotiropoulos, S. N. (2024). Denoising Diffusion MRI: Considerations and implications for analysis. Imaging Neuroscience, 2, 1–29.

Miller, K. L., & Pauly, J. M. (2003). Nonlinear phase correction for navigated diffusion imaging. Magnetic Resonance in Medicine: An Official Journal of the International Society for Magnetic Resonance in Medicine, 50(2), 343–353.

Miller, K. L., Stagg, C. J., Douaud, G., Jbabdi, S., Smith, S. M., Behrens, T. E., Jenkinson, M., Chance, S. A., Esiri, M. M., & Voets, N. L. (2011). Diffusion imaging of whole, post-mortem human brains on a clinical MRI scanner. NeuroImage, 57(1), 167–181.

Moeller, S., Pisharady, P. K., Ramanna, S., Lenglet, C., Wu, X., Dowdle, L., Yacoub, E., Uğurbil, K., & Akçakaya, M. (2021). NOise reduction with DIstribution Corrected (NORDIC) PCA in dMRI with complex-valued parameter-free locally low-rank processing. NeuroImage, 226, 117539.

Mugler III, J. P., & Brookeman, J. R. (1990). Three-dimensional magnetization-prepared rapid gradient-echo imaging (3D MP RAGE). Magnetic resonance in medicine, 15(1), 152–157.

Ouyang, M., Kang, H., Detre, J. A., Roberts, T. P., & Huang, H. (2017). Short-range connections in the developmental connectome during typical and atypical brain maturation. Neuroscience & Biobehavioral Reviews, 83, 109–122.

Pauly, J., Le Roux, P., Nishimura, D., & Macovski, A. (1991). Parameter relations for the Shinnar-Le Roux selective excitation pulse design algorithm (NMR imaging). IEEE transactions on medical imaging, 10(1), 53–65.

Polimeni, J. R., Bhat, H., Witzel, T., Benner, T., Feiweier, T., Inati, S. J., Renvall, V., Heberlein, K., & Wald, L. L. (2016). Reducing sensitivity losses due to respiration and motion in accelerated echo planar imaging by reordering the autocalibration data acquisition. Magnetic resonance in medicine, 75(2), 665–679.

Saleem, K. S., Avram, A. V., Glen, D., Yen, C. C.-C., Frank, Q. Y., Komlosh, M., & Basser, P. J. (2021). High-resolution mapping and digital atlas of subcortical regions in the macaque monkey based on matched MAP-MRI and histology. NeuroImage, 245, 118759.

Setsompop, K., Fan, Q., Stockmann, J., Bilgic, B., Huang, S., Cauley, S. F., Nummenmaa, A., Wang, F., Rathi, Y., & Witzel, T. (2018). High-resolution in vivo diffusion imaging of the human brain with generalized slice dithered enhanced resolution: Simultaneous multislice (g S lider-SMS). Magnetic resonance in medicine, 79(1), 141–151.

Smith, S. M., Jenkinson, M., Woolrich, M. W., Beckmann, C. F., Behrens, T. E., Johansen-Berg, H., Bannister, P. R., De Luca, M., Drobnjak, I., & Flitney, D. E. (2004). Advances in functional and structural MR image analysis and implementation as FSL. NeuroImage, 23, S208–S219.

Song, A. W., Chang, H.-C., Petty, C., Guidon, A., & Chen, N.-K. (2014). Improved delineation of short cortical association fibers and gray/white matter boundary using whole-brain three-dimensional diffusion tensor imaging at submillimeter spatial resolution. Brain connectivity, 4(9), 636–640.

Tendler, B. C., Hanayik, T., Ansorge, O., Bangerter-Christensen, S., Berns, G. S., Bertelsen, M. F., Bryant, K. L., Foxley, S., van den Heuvel, M. P., & Howard, A. F. (2022). The Digital Brain Bank, an open access platform for post-mortem imaging datasets. Elife, 11, e73153.

Tian, Q., Bilgic, B., Fan, Q., Liao, C., Ngamsombat, C., Hu, Y., Witzel, T., Setsompop, K., Polimeni, J. R., & Huang, S. Y. (2020). DeepDTI: High-fidelity six-direction diffusion tensor imaging using deep learning. NeuroImage, 219, 117017.

Tian, Q., Li, Z., Fan, Q., Polimeni, J. R., Bilgic, B., Salat, D. H., & Huang, S. Y. (2022). SDnDTI: Self-supervised deep learning-based denoising for diffusion tensor MRI. NeuroImage, 253, 119033.

Tournier, J.-D., Calamante, F., & Connelly, A. (2007). Robust determination of the fibre orientation distribution in diffusion MRI: non-negativity constrained super-resolved spherical deconvolution. NeuroImage, 35(4), 1459–1472.

Tournier, J.-D., Calamante, F., Gadian, D. G., & Connelly, A. (2004). Direct estimation of the fiber orientation density function from diffusion-weighted MRI data using spherical deconvolution. NeuroImage, 23(3), 1176–1185.

Tournier, J.-D., Smith, R., Raffelt, D., Tabbara, R., Dhollander, T., Pietsch, M., Christiaens, D., Jeurissen, B., Yeh, C.-H., & Connelly, A. (2019). MRtrix3: A fast, flexible and open software framework for medical image processing and visualisation. NeuroImage, 202, 116137.

Tournier, J. D., Calamante, F., & Connelly, A. (2010). Improved probabilistic streamlines tractography by 2nd order integration over fibre orientation distributions. Proceedings of the international society for magnetic resonance in medicine, Stockholm, Sweden.

Uecker, M., Lai, P., Murphy, M. J., Virtue, P., Elad, M., Pauly, J. M., Vasanawala, S. S., & Lustig, M. (2014). ESPIRiT—an eigenvalue approach to autocalibrating parallel MRI: where SENSE meets GRAPPA. Magnetic resonance in medicine, 71(3), 990–1001.

Veraart, J., Novikov, D. S., Christiaens, D., Ades-Aron, B., Sijbers, J., & Fieremans, E. (2016). Denoising of diffusion MRI using random matrix theory. NeuroImage, 142, 394–406.

Wang, F., Dong, Z., Tian, Q., Liao, C., Fan, Q., Hoge, W. S., Keil, B., Polimeni, J. R., Wald, L. L., & Huang, S. Y. (2021). In vivo human whole-brain Connectom diffusion MRI dataset at 760 µm isotropic resolution. Scientific Data, 8(1), 122.

Wu, W., Koopmans, P. J., Andersson, J. L., & Miller, K. L. (2019). Diffusion acceleration with Gaussian process estimated reconstruction (DAGER). Magnetic resonance in medicine, 82(1), 107–125.

Wu, W., Koopmans, P. J., Frost, R., & Miller, K. L. (2016). Reducing slab boundary artifacts in three-dimensional multislab diffusion MRI using nonlinear inversion for slab profile encoding (NPEN). Magnetic resonance in medicine, 76(4), 1183–1195.

Wu, W., Poser, B. A., Douaud, G., Frost, R., In, M.-H., Speck, O., Koopmans, P. J., & Miller, K. L. (2016). High-resolution diffusion MRI at 7T using a three-dimensional multi-slab acquisition. NeuroImage, 143, 1–14.

Zhang, T., Pauly, J. M., Vasanawala, S. S., & Lustig, M. (2013). Coil compression for accelerated imaging with Cartesian sampling. Magnetic resonance in medicine, 69(2), 571–582.

Zhang, Y., Brady, M., & Smith, S. (2001). Segmentation of brain MR images through a hidden Markov random field model and the expectation-maximization algorithm. IEEE transactions on medical imaging, 20(1), 45–57.

Zhu, S., Huszar, I. N., Cottaar, M., Daubney, G., Eichert, N., Hanayik, T., Khrapitchev, A. A., Mars, R. B., Mollink, J., & Sallet, J. (2024). Imaging the structural connectome with hybrid diffusion MRI-microscopy tractography. bioRxiv, 2024.2001. 2008.574641.

